# Multifunctional properties of Nej1^XLF^ C-terminus promote end-joining and impact DNA double-strand break repair pathway choice

**DOI:** 10.1101/2022.01.13.476264

**Authors:** Aditya Mojumdar, Nancy Adam, Jennifer Cobb

## Abstract

A DNA double strand break (DSB) is primarily repaired by one of two canonical pathways, non-homologous end-joining (NHEJ) and homologous recombination (HR). NHEJ requires no or minimal end processing for ligation, whereas HR requires 5’ end resection followed by a search for homology. The main event that determines the mode of repair is the initiation of 5’ resection because if resection starts, then NHEJ cannot occur. Nej1 is a canonical NHEJ factor that functions at the cross-roads of repair pathway choice and prior to its function in stimulating Dnl4 ligase. Nej1 competes with Dna2, inhibiting its recruitment to DSBs and thereby inhibiting resection. The highly conserved C-terminal region (CTR) of Nej1 (330-338) is important for two events that drive NHEJ, stimulating ligation and inhibiting resection, but it is dispensable for end-bridging. By combining *nej1* point mutants with nuclease-dead *dna2*-1, we find that Nej1-F335 is essential for end-joining whereas V338 promotes NHEJ indirectly through inhibiting Dna2-mediated resection.

**Highlights:**

- Nej1 C-terminus is critical for repair pathway choice.
- The KKRK region of Nej1 is important for interactions with ssDNA and dsDNA.
- Nej1-F335 and V338 are key residues for end-joining and inhibition of resection at DSB.
- Nej1-mediated end-bridging is not sufficient for end-joining repair.

## 1. INTRODUCTION

The two major pathways of DNA double strand break (DSB) repair are non-homologous end-joining (NHEJ) and homologous recombination (HR). Although HR is the predominant pathway in *S. cerevisiae* (budding yeast), core NHEJ components are highly conserved from yeast to humans [1, 2]. Repair factors have largely been defined for their involvement in either HR or NHEJ, however an individual component can drive repair events in multiple ways. Many of the repair factors function at the cross-roads of these two canonical pathways, impacting whether DNA ends at the DSB will be ligated or resected.

Once a double strand break occurs, yKu70/80 (Ku) and Mre11-Rad50-Xrs2 (MRX) localize to the site of damage independently of each other. Nej1 was first identified as a NHEJ factor and is recruited to DSBs through interactions with yKu70/80 (Ku), but more recently it was also shown to be recruited through interactions with Mre11 [3]. Ku has a high affinity towards double stranded DNA ends and binds to broken DNA ends primarily to protect them from nucleolytic degradation. Subsequently, the other core NHEJ factors, Lif1-Dnl4 and Nej1, get recruited [4]. Dnl4 ligase, together with Lif1 and stimulated by Nej1, completes repair by ligating the DNA ends [5-8].

Nej1 also promotes NHEJ indirectly by downregulating HR through at least two different mechanisms. First, Nej1 stabilizes Ku bound at DNA ends as a way to prevent resection [9, 10]. However, in *nej1*Δ mutants increased 5’ resection does not depend on Exo1 [11], therefore Ku likely continues to provide a level of DNA end protection when *NEJ1* is deleted as Exo1 is the preferred nuclease for resection in the absence of Ku [12]. Secondly, Nej1 inhibits 5’ resection by competing with Dna2 at the break site [3, 13]. Nej1 has a competitive and antagonistic relationship with Dna2 for binding to Sgs1 and Mre11. Dna2 nuclease resects DNA ends at a DSB in addition to its essential role in replication fork processing and Okazaki fragment maturation [13-19]. When Mre11 nuclease fails to initiate resection by loss of its intrinsic activity or by SAE2 deletion, Dna2-Sgs1 can compensate. Our previous work added to this understanding as we showed that Nej1 regulates resection and Dna2 recruitment even in *MRE11*^+^ or otherwise wild type cells, and that the presence of Nej1 prevents large deletions for developing during DSB repair [3, 13]. In addition to stimulating Dnl4 ligase and inhibiting Dna2 nuclease, Nej1 also functions in DNA end-bridging. This role is not shared with the NHEJ factors Ku and Dnl4 or the 5’ resection factors Exo1 and Sgs1. However, the MRX complex is important for bridging. The structural features of the complex, rather than Mre11 nuclease activity, are important for end-bridging [3, 13].

The N-terminus of Nej1 interacts with Ku and the C-terminus of Nej1 (173-342aa) interacts with Lif1 [20, 21]. The regions in Nej1 that interact with Ku and Lif1 are reversed from their counterparts in humans as the N-terminus of XLF (human homologue of Nej1) interacts with XRCC4 (Lif1) and the C-terminus of XLF interacts with Ku [20, 22-24]. The C-terminus of Nej1 is essential for NHEJ repair. KKRK (330-334 aa) bind DNA *in vitro* and serves as a nuclear localization signal (NLS), whereas FGKV (335-338 aa) interacts with Lif1 [21, 25]. The sequence conservation of this region among Nej1 and XLF homologues in various organisms makes the C-terminus an interesting target for detailed molecular investigation [21, 23].

In general, there is an inverse relationship between 5’ resection and NHEJ repair. Here we investigated whether the C-terminal region of Nej1 is important for bridging DNA ends and for inhibiting Dna2. We utilized nuclease dead *dna2*-1 as a genetic tool in combination with *nej1* mutants to identify key residues in the C-terminal region of Nej1 that are important for end-joining directly versus those that drive NHEJ indirectly by way of inhibiting Dna2. Our work showed that F335 was critical for NHEJ independently of resection, whereas V338 promoted NHEJ primarily by inhibiting resection.

## 2. MATERIALS AND METHODS

### 2.1 Media details

All the yeast strains used in this study are listed in Table S1 and were obtained by crosses. The strains were grown on various media in experiments described below. For HO-induction of a DSB, YPLG media is used (1% yeast extract, 2% bactopeptone, 2% lactic acid, 3% glycerol and 0.05% glucose). For the continuous DSB assay, YPA plates are used (1% yeast extract, 2% bacto peptone, 0.0025% adenine) supplemented with either 2% glucose or 2% galactose. For the mating type assays, YPAD plates are used (1% yeast extract, 2% bacto peptone, 0.0025% adenine, 2% dextrose).

### 2.2 Chromatin Immunoprecipitation

ChIP assay was performed as described previously [3]. Cells were cultured overnight in YPLG at 25°C. Cells were then diluted to equal levels (5 × 10^6^ cells/ml) and were cultured to one doubling (3-4 hrs) at 30°C. 2% GAL was added to the YPLG and cells were harvested and crosslinked at various time points using 3.7% formaldehyde solution. Following crosslinking, the cells were washed with ice cold PBS and the pellet stored at -80°C. The pellet was re-suspended in lysis buffer (50mM Hepes pH 7.5, 1mM EDTA, 80mM NaCl, 1% Triton, 1mM PMSF and protease inhibitor cocktail) and cells were lysed using Zirconia beads and a bead beater. Chromatin fractionation was performed to enhance the chromatin bound nuclear fraction by spinning the cell lysate at 13,200rpm for 15 minutes and discarding the supernatant. The pellet was re-suspended in lysis buffer and sonicated to yield DNA fragments (∼500bps in length). The sonicated lysate was then incubated in beads + anti-HA/Myc/FLAG Antibody or unconjugated beads (control) for 2 hrs at 4°C. The beads were washed using wash buffer (100mM Tris pH 8, 250mM LiCl, 150mM (HA/Flag Ab) or 500mM (Myc Ab) NaCl, 0.5% NP-40, 1mM EDTA, 1mM PMSF and protease inhibitor cocktail) and protein-DNA complex was eluted by reverse crosslinking using 1%SDS in TE buffer, followed by proteinase K treatment and DNA isolation via phenol-chloroform-isoamyl alcohol extraction. Quantitative PCR was performed using the Applied Biosystem QuantStudio 6 Flex machine. PerfeCTa qPCR SuperMix, ROX was used to visualize enrichment at HO2 (0.5kb from DSB) and SMC2 was used as an internal control.

### 2.3 Continuous DSB assay and identification of mutations in survivors

Cells were grown overnight in YPLG media at 25°C to saturation. Cells were collected by centrifugation at 2500rpm for 3 minutes and pellets were washed 1x in ddH_2_O and re-suspended in ddH_2_O. Cells were counted and spread on YPA plates supplemented with either 2% GLU or 2% GAL. On the Glucose plates 1×10^3^ total cells were added and on the galactose plates 1×10^5^ total cells were added. The cells were incubated for 3-4 days at room temperature and colonies counted on each plate. Survival was determined by normalizing the number of surviving colonies in the GAL plates to number of colonies in the GLU plates. 100 survivors from each strain were scored for the mating type assay as previously described [11].

### 2.4 qPCR based Ligation Assay

Cells from each strain were grown overnight in 15ml YPLG to reach an exponentially growing culture of 1×10^7^ cells/mL. Next, 2.5mL of the cells were pelleted as ‘No break’ sample, and 2% GAL was added to the remaining cells, to induce a DSB. 2.5ml of cells were pelleted after 3hr incubation as timepoint 0 sample. Following that, GAL was washed off and the cells were released in YPAD and respective timepoint samples were collected. Genomic DNA was purified using standard genomic preparation method by isopropanol precipitation and ethanol washing, and DNA was re-suspended in 100uL ddH_2_O. Quantitative PCR was performed using the Applied Biosystem QuantStudio 6 Flex machine. PowerUp SYBR Green Master Mix was used to quantify resection at HO6 (at DSB) locus. Pre1 was used as an internal gene control for normalization. Signals from the HO6/Pre1 timepoints were normalized to ‘No break’ signals and % Ligation was determined. The primer sequences are listed in Table S3.

### 2.5 qPCR based Resection Assay

Cells from each strain were grown overnight in 15ml YPLG to reach an exponentially growing culture of 1×10^7^ cells/mL. Next, 2.5mL of the cells were pelleted as timepoint 0 sample, and 2% GAL was added to the remaining cells, to induce a DSB. Following that, respective timepoint samples were collected. Genomic DNA was purified using standard genomic preparation method by isopropanol precipitation and ethanol washing, and DNA was re-suspended in 100mL ddH_2_O. Genomic DNA was treated with 0.005μg/μL RNase A for 45min at 37°C. 2μL of DNA was added to tubes containing CutSmart buffer with or without RsaI restriction enzyme and incubated at 37°C for 2hrs. Quantitative PCR was performed using the Applied Biosystem QuantStudio 6 Flex machine. PowerUp SYBR Green Master Mix was used to quantify resection at MAT1 (0.15kb from DSB) locus. Pre1 was used as a negative control. RsaI cut DNA was normalized to uncut DNA as previously described to quantify the %ssDNA [26]. The primer sequences are listed in Table S3.

### 2.6 Microscopy of end-bridging at DSBs

Cells derived from the parent strain JC-4066 were diluted and grown overnight in YPLG at 25°C to reach a concentration of 1×10^7^ cells/ml. Cells were treated with 2% GAL for 2 hours and cell pellets were collected and washed 2 times with PBS. After the final wash, cells were placed on cover slips and imaged using a fully motorized Nikon Ti Eclipse inverted epi-fluorescence microscope. Z-stack images were acquired with 200 nm increments along the z plane, using a 60X oil immersion 1.4 N.A. objective. Images were captured with a Hamamatsu Orca flash 4.0 v2 sCMOS 16-bit camera and the system was controlled by Nikon NIS-Element Imaging Software (Version 5.00). All images were deconvolved with Huygens Essential version 18.10 (Scientific Volume Imaging, The Netherlands, http://svi.nl), using the Classic Maximum Likelihood Estimation (CMLE) algorithm, with SNR:40 and 50 iterations. To measure the distance between the GFP and mCherry foci, the ImageJ plug-in Distance Analysis (DiAna) was used [27]. Distance measurements represent the shortest distance between the brightest pixel in the mCherry channel and the GFP channel. Each cell was measured individually and > 50 cells were analyzed per condition per biological replicate.

### 2.7 Protein Expression and Purification

The plasmids containing the gene used for protein expression are listed in Table S2. The plasmids were transformed in wild type yeast cells (JC-727), they were grown overnight at 30°C in -TRP 2% GLU liquid media. The following day cells were pelleted and washed thrice with autoclaved ddH_2_O and released in -TRP 2% GAL liquid media. The wild-type and mutant proteins were expressed under GAL at 25°C for 24 hrs. The cells were harvested and lysed in lysis buffer (50mM Hepes pH 7.5, 1mM EDTA, 80mM NaCl, 1% Triton, 1mM PMSF, protease inhibitor cocktail and DNaseI) using Zirconia beads and a bead beater. The lysate was then incubated with beads conjugated with anti-HA Antibody for 2 hrs at 4°C. The beads were washed using wash buffer (100mM Tris pH 8, 250mM LiCl, 150mM NaCl, 0.5% NP-40, 1mM EDTA, 1mM PMSF and protease inhibitor cocktail) and protein was eluted using 100mM Glycine pH 2.0 and the eluted protein was immediately collected in 10X TBS buffer to avoid aggregation. Protein concentration was determined by measuring the absorbance at 280 nm, and protein purity was analyzed on SDS-PAGE and WB.

### 2.8 ThermoFluor Assay

To determine the protein stability and to compare the various mutants, a heat denaturation analysis was carried out using a CFX96 Touch real time quantitative PCR system (Bio-Rad) [28]. Protein stability measurements were performed in a buffer containing 20 mm Tris, pH 8.0, 250 mm NaCl, 5% glycerol (v/v), and 5 mm β-mercaptoethanol. The 25 μl of reaction mix contained 5 μm protein and SYPRO Orange (Invitrogen). Heat denaturing curves were observed within the temperature range of 37–85 °C, at a ramp rate of 1.8 °C/min, collecting data every 10 s. The data obtained was normalized using GraphPad Prism software.

### 2.9 DNA binding assay

The fluorescence polarization assay was performed by incubating 10 μL solution consisting of 10nM FAM labeled probes (ss and dsDNA) and final protein concentrations ranging from 0 to 4uM in 10 mM Tris-HCL pH 7.0 at room temperature for 10 min. The fluorescence polarization of the resulting solutions was measured by using PHERAstar FSX plate reader from BMG Labtech using FP 485 520 520 filter. The oligo sequences are listed in Table S3.

### 2.10 Western Blot

Cells were lysed by re-suspending them in lysis buffer (with PMSF and protease inhibitor cocktail tablets) followed by bead beating. The protein concentration of the whole cell extract was determined using the NanoDrop. Equal amounts of whole cell extract were added to wells of 10% polyacrylamide SDS gel followed by transfer to Nitrocellulose membranes at 100V for 80mins. The membrane was Ponceau stained (which served as a loading control), followed by blocking in 10% milk-PBST for 1hour at room temperature. Respective primary antibody solution (1:1000 dilution) was added and left for overnight incubation at 4°C. The membranes were then washed with PBST and left for 1 hour with secondary antibody (1:10000), followed by washing and adding ECL substrates prior to imaging.

## 3. RESULTS

### 3.1 Nej1 C-terminus is essential for end-ligation

The C-terminus of Nej1 is essential for NHEJ. This region binds DNA, contains a nuclear localization signal, KKRK (331-334), and interacts with Lif1 through FGVK (335-338) (Fig. 1A) [21, 25]. We previously demonstrated that when SV40 –NLS was fused to the N-terminus of *nej1Δ*C (1-330aa) and *nej1*-1 (KKRK→ AAAA) mutants that nuclear localization was restored. However, consistent with our earlier work, the survival of both NLS fusion mutants and *nej1*-2 (FGKV→ AAAA) was markedly reduced after a DSB was created in the inducible HO-DSB system (Fig. S1A, [21]).

**Figure 1.**
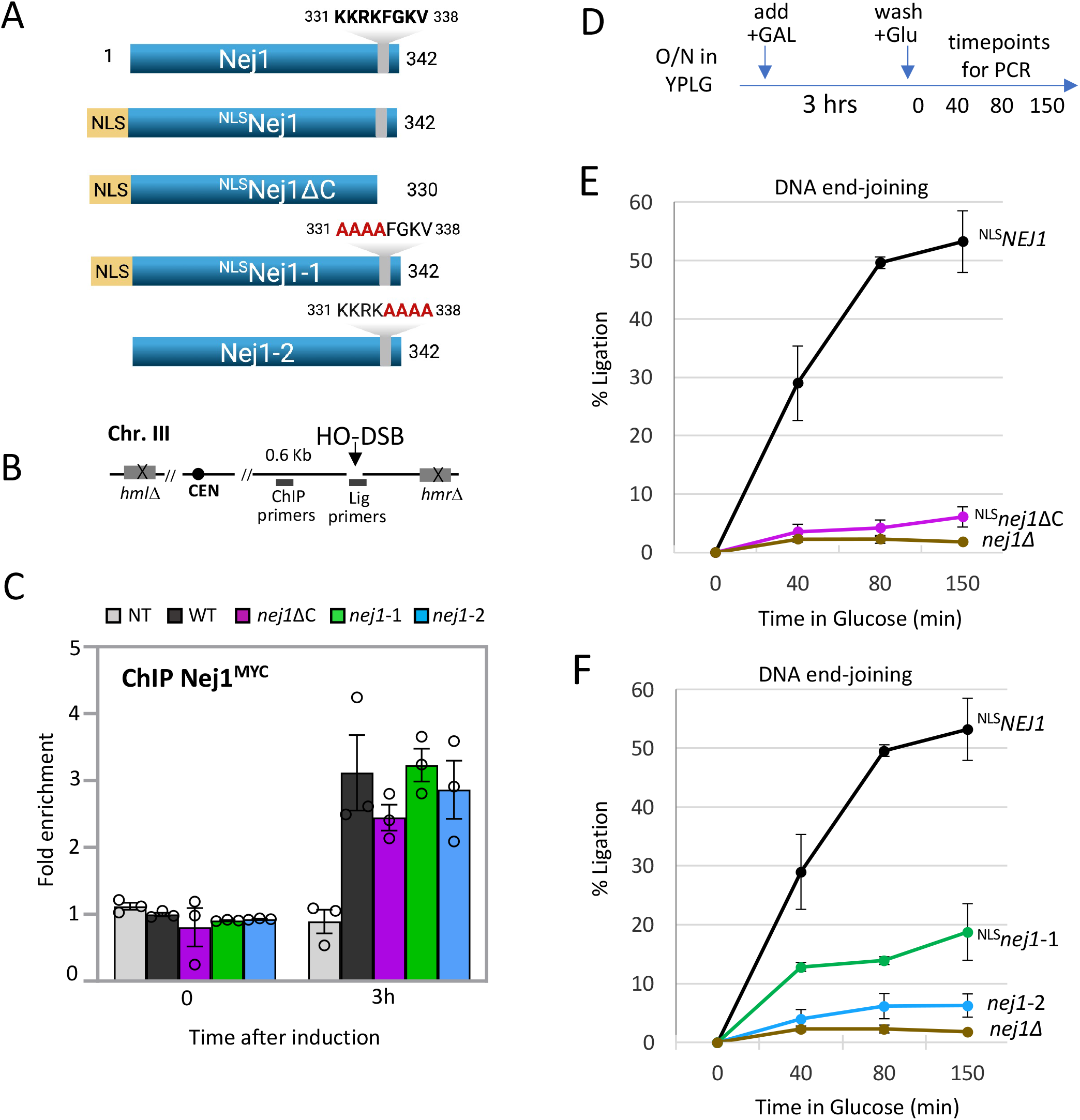
Nej1 C-terminus is essential for end-ligation stimulation. **(A)** Schematic representation of Nej1 C-terminal mutants used in this study. **(B)** Schematic representation of regions around the HO cut site on chromosome III. The ChIP probe used in this study is 0.6kb from the DSB. Primers at HO cut site is used in qPCR ligation assays. **(C)** Enrichment of Nej1^Myc^ at DSB, at 0 and 3 hours under DSB, in wild type *NEJ1*^*+*^ (JC-3193), *nej1*ΔC (JC-3194), *nej1*-1 (JC-3215), *nej1*-2 (JC-3134) and no tag control (JC-727) was determined. The fold enrichment is normalized to recovery at the SMC2 locus. **(D)** Schematic representation of the ligation assay set up. Cells were grown overnight in YPLG to a certain number, galactose was added to induce DSB for 3 hours, followed by washing the cells and releasing them in glucose rich media (YPAD) to recover from DSB. Time points were collected and genomic DNA was extracted that was later used for qPCR using the respective probes at HO cut site. **(E and F)** qPCR-based ligation assay of DNA at HO DSB, as measured by % Ligation, at 0, 40, 80 and 150 min in cycling cells in glucose post DSB. Strains used were wild type *NEJ1*^*+*^ (JC-3193), *nej1*Δ (JC-1342), *nej1*ΔC (JC-3194), *nej1*-1 (JC-3215) and *nej1*-2 (JC-3134). For all the experiments - error bars represent the standard error of three replicates.

To generate a map for how these C-terminal residues impact DSB repair more directly we measured protein localization and end-joining capabilities of the three *nej1* mutants *in vivo* at the HO-DSB (Fig. 1B). ChIP was performed with Myc-tagged Nej1 and showed that all three mutants were recruited to the break site at levels similarly to wild type (Fig. 1C). Residues 335-338 in the C-terminus of Nej1 interacts with Lif1 [21], however the recruitment of Lif1, and the other NHEJ factors Ku70 and Dnl4, were not significantly affected in these *nej1* mutants (Fig. S1B-D).

We next measure the ability of these mutants to ligate a DSB *in vivo*. Rejoining of DNA ends was determined in cells where the DSB was induced by galactose for 3 hours, before washing the cells and releasing them into glucose to prevent further re-cutting (Fig. 1D). At the indicated time-points, genome DNA was prepared followed by quantitative PCR with primers flanking the break site (Fig. 1B, Lig primers), which would amplify only if the broken DNA ends joined. End-joining in ^NLS^*nej1Δ*C and *nej1*-2 were indistinguishable from *nej1Δ* mutants (Fig. 1E andF), and and in ^NLS^*nej1*-1 it was reduced to ∼ 35% of ^NLS^*NEJ1* (Fig. 1F). Cells expressing ^NLS^Nej1 showed similar ligation rates to wild type cells (Fig. S1E and F).

Nej1, but not Dnl4 or Ku, is important for DNA end-bridging, therefore NHEJ defects could be partly due to compromised end-bridging in the C-terminal mutants. We determined whether end-bridging was intact by measuring the distance across the DSB in cells where the TetO and LacO arrays were integrated 3.2 Kb upstream 5.2Kb downstream from the HO cut site respectively. Expression of GFP-tagged TetR and mCherry-tagged LacO enabled us to visualize the two sides of the break site by fluorescence microscopy (Fig. S1F). Consistent with our earlier work, end-bridging was disrupted in cells where *NEJ1* was deleted with a ∼ 2-fold increase inthe distance between the fluorophores compared to wild type. Both ^NLS^*nej1Δ*C and ^NLS^*nej1*-1 mutants showed a very modest, but non-significant, end-bridging defect and there was no defect seen in *nej1*-2 mutants (Fig. 1F). Taken together, decreased ligation in these C-terminal mutants was not caused by their failure to recruit nor disruptions in end-bridging.

### 3.2 The C-terminal ‘KKRK’ motif binds ssDNA and dsDNA

In addition to serving as a nuclear localization signal, the ‘KKRK’ motif of Nej1 was previously shown to bind 20-300 bp of dsDNA [25]. Indeed, defects in DNA binding could contribute to defects in end-joining, and it was not known whether the FGKV region was important for DNA binding. To this end, we overexpressed and purified Nej1 proteins, full-length and C-terminal mutants, and measured their affinity to single-stranded DNA (ssDNA; S1) and double-stranded (dsDNA; S1:S2), two potential Nej1 substrates at DSBs.

We expressed HA-tagged Nej1 proteins in yeast from Y2H vectors induced overnight with galactose and purified with αHA-beads (Fig. 3A). The purified proteins were predicted to fold properly based on a thermal shift assay [28]. In this thermofluor assay, SYPRO orange dye is added to the protein sample, which is further subjected to a gradual increase in temperature leading to the denaturation of the protein. As the hydrophobic residues of the protein get exposed, the dye binds to these residues producing a fluorescence signal. All the Nej1 C-terminal mutant proteins exhibited a behavior similar to the WT protein, confirming that the overall protein folding was not affected due to the mutations (Fig. 3B). This was not unexpected given the C-terminus of Nej1 has been predicted to be a high-mobility unstructured region and not critical for overall architecture [29].

The purified proteins were used to compare binding affinities of full-length Nej1 and mutants with single-stranded DNA (ssDNA; S1) and double-stranded (dsDNA; S1:S2). Fluorescently labeled probes (Table S3; [30]) were incubated with increasing protein concentrations (0-4 μM), and protein-DNA binding was determined by fluorescence polarization. Purified full length Nej1 showed binding to both dsDNA and ssDNA (Fig. 2C-D). Purified Nej1-2 interacted with dsDNA and ssDNA similarly to full length Nej1 (Fig. 2C and 2D). However, Nej1ΔC and Nej1-1 showed reduced affinity for DNA, with binding to ssDNA being highly compromised (Fig. 2D). This is notable given HO-DSBs have complimentary ssDNA overhangs that are compatible for direct ligation by NHEJ [31].

**Figure 2.**
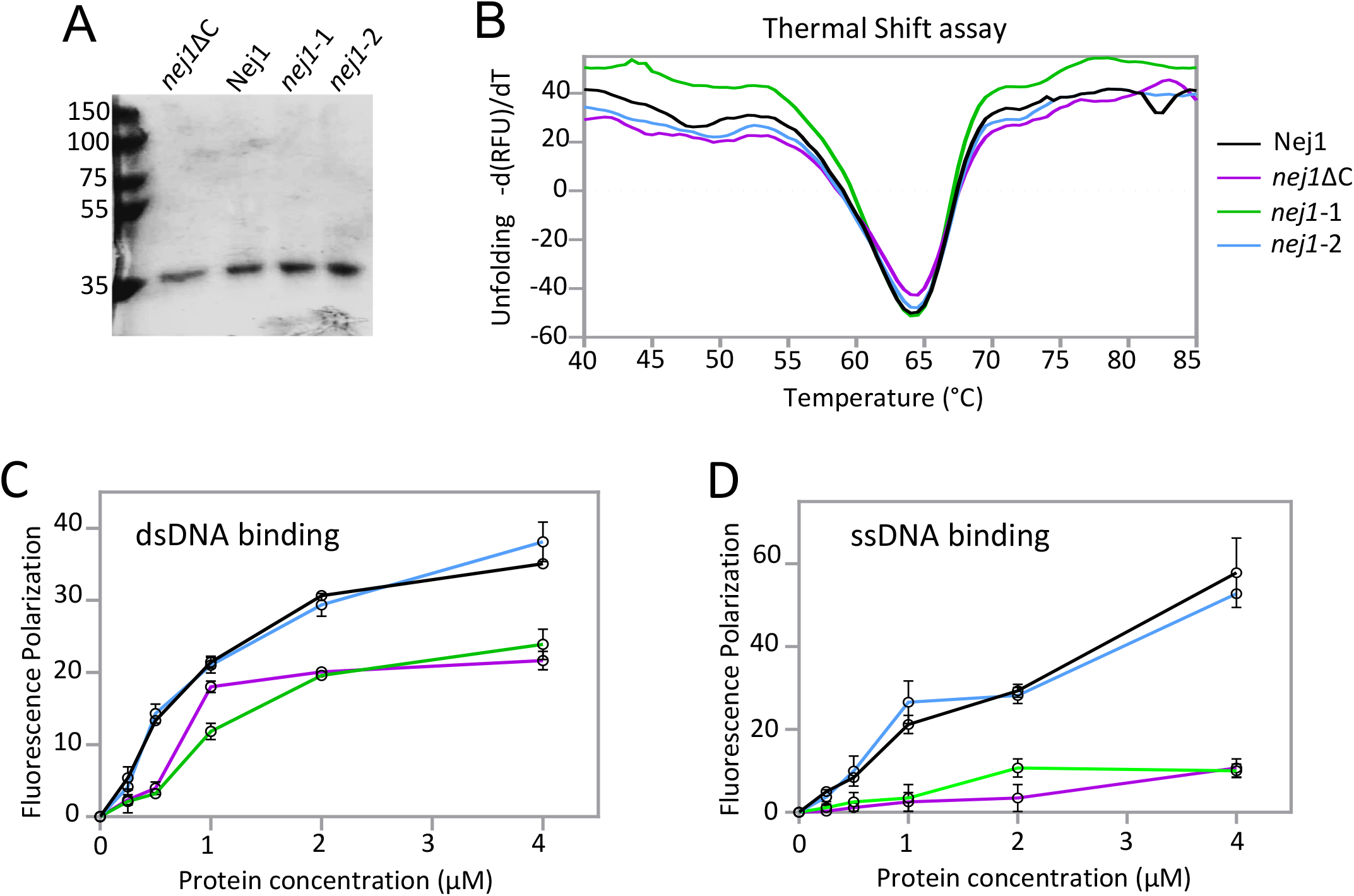
The C-terminal ‘KKRK’ motif facilitates end-ligation by binding DNA overhangs at DSB. **(A)** SDS-PAGE gel of purified proteins – Nej1 (WT), *nej1*ΔC, *nej1*-1 and *nej1*-2. **(B)** Heat denaturation profiles of Nej1 variants. The curves depicting the unfolded proteins in terms of −(dRFU)/dT. **(C and D)** DNA binding activity was measured using Fluorescence polarization method. The protein concentration was increased from 0-4uM. The plots depict the comparison of DNA binding activity of Nej1 variants on two substrates: **(C)** dsDNA (S1:S2) and **(D)** ssDNA (S1). The error bars represent the standard error of three replicates.

Given these results, one potential role for Nej1 in facilitating end-ligation might be in stabilizing ssDNA overhangs at the DSB through interaction with its ‘KKRK’ motif. Nevertheless, the data presented thus far help clarify two aspects of Nej1 biology. First, the ability of Nej1 to localize to DSBs is independent of DNA binding as all mutants, including *nej1*ΔC and *nej1*-1, localize to the DSB *in vivo* (likely through interactions with Ku which map to the N-terminus of Nej1).

Secondly, the ability of Nej1 to promote end-ligation is distinct of its ability to bind DNA. End-joining did not proceed in *nej1*-2 mutant cells, even when Nej1-2 and the other NHEJ factors localize (Fig 1 and S), however, *in vitro* Nej1-2 interactions with DNA did not diminish.

### 3.3 Nej1 C-terminal region inhibits Dna2 at DSB

As Nej1 inhibits resection, decreased end-ligation might be an indirect effect of increased 5’ resection in the Nej1 C-terminal mutants. As previously described, resection was determined in cells upon DSB induction by the addition of galactose at the indicated timepoints from 0 to 150 mins. ([13]; Fig. 3A). Genomic DNA was prepared and digested with the *Rsa*I restriction enzyme followed by quantitative PCR. This method relies on a *Rsa*I cut site located 0.15 Kb from the DSB (Fig. 3A). If resection progresses past this recognition site, then single stranded DNA will be generated and the site would not be cleaved by *Rsa*I. The region can be amplified by PCR [3, 11, 26, 32]. Consistent with earlier work, deletion of *NEJ1* resulted in hyper 5’ resection. This phenotype was observed in both *nej1*-2 and ^NLS^*nej1Δ*C mutants, however resection in ^NLS^*nej1*-1 was similar to *NEJ1+* cells (Fig. 3B). Previously we showed that Nej1 inhibits end-resection by inhibiting Dna2 nuclease recruitment to DSB. We measured Dna2 recruitment by performing ChIP [3, 13]. In line with increased resection, Dna2 recovery by ChIP increased in ^NLS^*nej1*ΔC and *nej1*-2, but not ^NLS^*nej1*-1 mutants (Fig. 3C).

**Figure 3.**
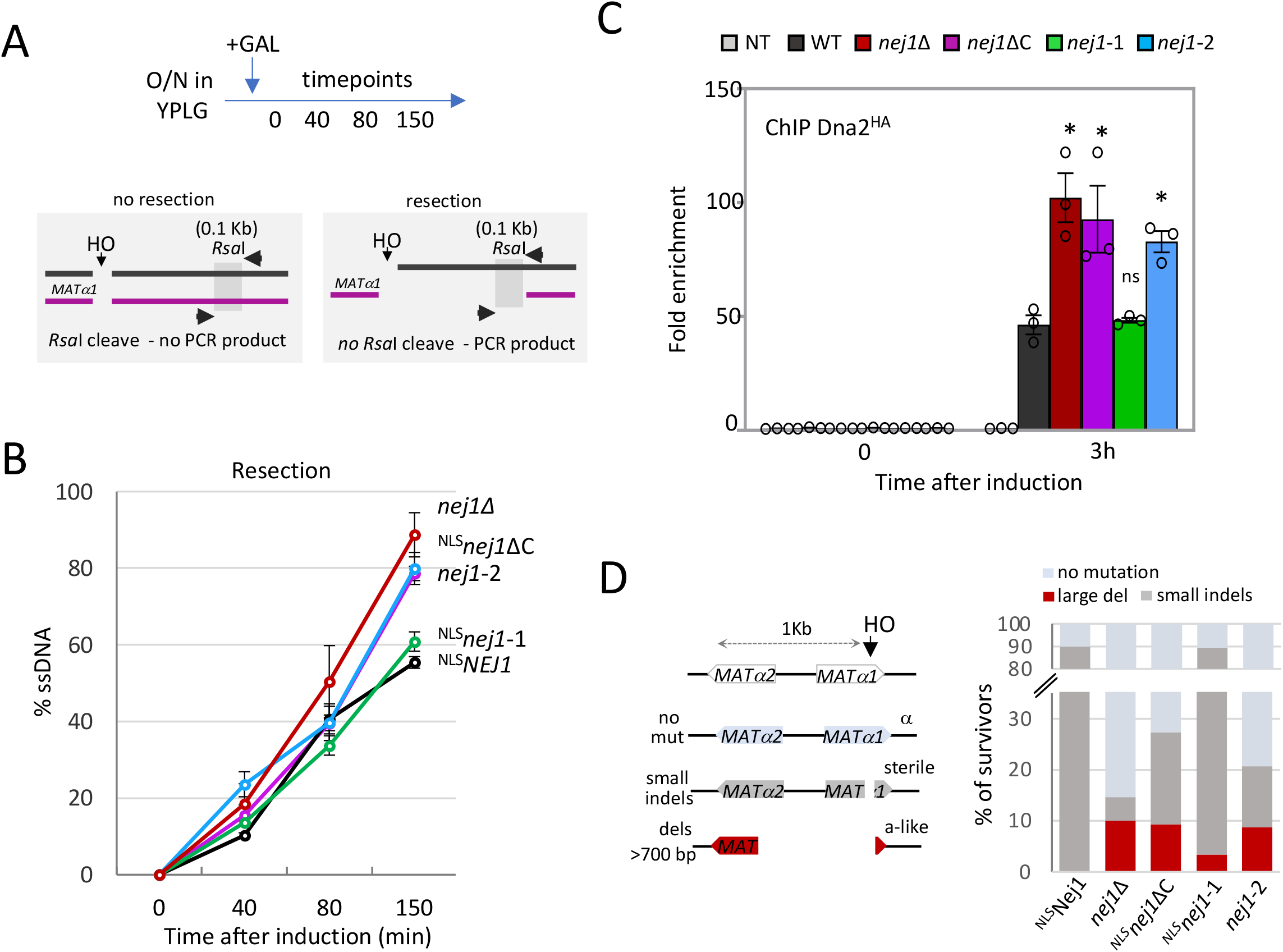
Nej1 C-terminal region inhibits Dna2 at DSB. **(A) Above:** Schematic representation of the resection assay set up. Cells were grown overnight in YPLG to a certain number, galactose was added to induce DSB. Time points were collected and genomic DNA was extracted that was later used for qPCR using the respective probes at RsaI cut site. **Below:** Schematic representation of the RsaI sites used in the qPCR resection assays, 0.15kb from the DSB. **(B)** qPCR-based resection assay of DNA 0.15kb away from the HO DSB, as measured by % ssDNA, at 0, 40, 80 and 150 min post DSB induction in cycling cells in wild type *NEJ1*^*+*^ (JC-3193), *nej1*Δ (JC-1342), *nej1*ΔC (JC-3194), *nej1*-1 (JC-3215) and *nej1*-2 (JC-3134). **(C)** Enrichment of Dna2^HA^ at 0.6kb from DSB 0 hour (no DSB induction) and 3 hours after DSB induction in wild type (JC-4117), *nej1*ΔC (JC-5866), *nej1*-1 (JC-5868), *nej1*-2 (JC-5867) and no tag control (JC-727) was determined. The fold enrichment is normalized to recovery at the SMC2 locus. **(D) Left:** Schematic representation of mating type analysis of survivors from persistent DSB induction assays. The mating phenotype is a read out for the type of repair: α survivors (light gray), sterile survivors (small insertions and deletions, dark gray) and “a-like” survivors (>700 bps deletion, red). **Right:** Mating type analysis of survivors from persistent DSB induction assays mentioned in (B). For all the experiments - error bars represent the standard error of three replicates. Significance was determined using 1-tailed, unpaired Student’s t test. All strains compared are marked (P<0.05*; P<0.01**; P<0.001***) unless specified with a line, strains are compared to WT.

Increased Dna2 recruitment and hyper-resection at DSB correlates with the development of large genomic deletion [3]. Therefore, to determine whether genomic alterations arise during DSB repair in a physiological system, we determined the mating type of survivors during continuous galactose exposure. The induced HO -DSB site is adjacent to two genes MATα1 and MATα2, that regulate mating type by activating alpha-type genes and inhibiting a-type genes (Fig. 3D). During DSB repair process, large deletions (>700 bp) that disrupt both α1 and α2 genes result in ‘a-like’ survivors (red), whereas small deletions or insertions lead to sterile survivors (gray). Both ^NLS^*nej1*ΔC and *nej1*-2 mutants showed a similar accumulation of a-like survivors to the level seen when *NEJ1* was deleted. However, large genomic deletions formed at a lower rate in ^NLS^*nej1*-1 survivors (Fig. 3D and Table 1).

**Table 1.** Survival and percentage of large deletions during continuous DSB.

| Genotype | Survival | std Dev<br>(+/-) | Survival relative<br>to WT (%) | Large<br>deletions (%) |
| --- | --- | --- | --- | --- |
| WT | $2.9 \times 10^{-3}$ | $5.5 \times 10^{-4}$ | 100.00% | 1 |
| *NLS**Nej1*<sup>+</sup> | $3.1 \times 10^{-3}$ | $4.9 \times 10^{-4}$ | 106.82% | 0 |
| *nej1*Δ | $1.7 \times 10^{-5}$ | $5.8 \times 10^{-6}$ | 0.57% | 10 |
| *NLS**nej1* (1-330) | $5.0 \times 10^{-5}$ | $2.7 \times 10^{-5}$ | 1.70% | 9 |
| *NLS**nej1-1* | $2.2 \times 10^{-4}$ | $3.5 \times 10^{-5}$ | 7.39% | 2 |
| *nej1-2* | $5.0 \times 10^{-5}$ | $2.0 \times 10^{-5}$ | 1.70% | 9 |
| *nej1*<sup>F335A</sup> | $5.3 \times 10^{-5}$ | $2.1 \times 10^{-5}$ | 1.82% | 8 |
| *nej1*<sup>V338A</sup> | $1.8 \times 10^{-4}$ | $3.5 \times 10^{-5}$ | 6.25% | 2 |
| *dna2-1* | $7.3 \times 10^{-3}$ | $1.3 \times 10^{-3}$ | 250.34% | 0 |
| *dna2-1* *NLS**Nej1*<sup>+</sup> | $6.8 \times 10^{-3}$ | $5.6 \times 10^{-4}$ | 234.66% | 0 |
| *dna2-1 nej1*Δ | $2.7 \times 10^{-5}$ | $5.8 \times 10^{-6}$ | 0.91% | 0 |
| *dna2-1* *NLS**nej1* (1-330) | $5.0 \times 10^{-5}$ | $2.0 \times 10^{-5}$ | 1.70% | 0 |
| *dna2-1* *NLS**nej1-1* | $1.1 \times 10^{-3}$ | $1.6 \times 10^{-4}$ | 36.82% | 0 |
| *dna2-1 nej1-2* | $5.3 \times 10^{-5}$ | $2.5 \times 10^{-5}$ | 1.82% | 0 |
| *dna2-1 nej1*<sup>F335A</sup> | $7.7 \times 10^{-5}$ | $1.5 \times 10^{-5}$ | 2.61% | 0 |
| *dna2-1 nej1*<sup>V338A</sup> | $2.0 \times 10^{-3}$ | $2.1 \times 10^{-4}$ | 68.52% | 0 |
| *ku70*Δ | $2.0 \times 10^{-5}$ | $1.0 \times 10^{-5}$ | 0.68% | 14 |
| *lif1*Δ | $4.7 \times 10^{-5}$ | $1.5 \times 10^{-5}$ | 1.59% | 9 |
| *dnl4*Δ | $4.7 \times 10^{-5}$ | $2.0 \times 10^{-5}$ | 1.59% | 12 |
| *dna2-1 ku70*Δ | $3.3 \times 10^{-5}$ | $1.5 \times 10^{-5}$ | 1.14% | 10 |
| *dna2-1 lif1*Δ | $3.7 \times 10^{-5}$ | $1.5 \times 10^{-5}$ | 1.25% | 0 |
| *dna2-1 dnl4*Δ | $4.0 \times 10^{-5}$ | $1.0 \times 10^{-5}$ | 1.36% | 0 |
| *sgs1*Δ | $3.8 \times 10^{-3}$ | $1.5 \times 10^{-3}$ | 130.68% | 0 |
| *mre11-3* | $3.8 \times 10^{-3}$ | $7.6 \times 10^{-4}$ | 130.68% | 0 |
| *exo1*Δ | $3.7 \times 10^{-3}$ | $1.0 \times 10^{-3}$ | 127.39% | 0 |
| *sae2*Δ | $5.6 \times 10^{-3}$ | $1.8 \times 10^{-3}$ | 189.77% | 0 |

The FGKV region was important for inhibiting Dna2 and 5’ resection, however this region was previously shown to be important for Lif1 binding. Therefore, we next wanted to investigate whether the residues stimulating ligation and Lif1 binding were also important in Dna2 inhibition.

### 3.4 Nej1 Phenylalanine-335 and Valine-338 promote NHEJ by inhibiting Dna2

Of the four amino acids in FGKV, Phenylalanine-335 (F335) and Valine-338 (V338) have a major role in Nej1-Lif1 binding and mutations in these residues led to reduced survival after HO induction (Fig. 4A; Table 1; [21]). Consistently, here we show that cells harbouring *nej1*^F335A^ and *nej1*^V338A^ were also defective in end-joining (Fig. 4B). The ∼3-fold reduction in *nej1*^V338A^ was somewhat surprising given that mutant could support ligation in a *Xho*I cut plasmid repair assay (Table 1; [21]). The defect in end-joining was not because of compromised end-bridging as both the mutants showed wild type level of end-bridging (Fig. S2A).

**Figure 4.**
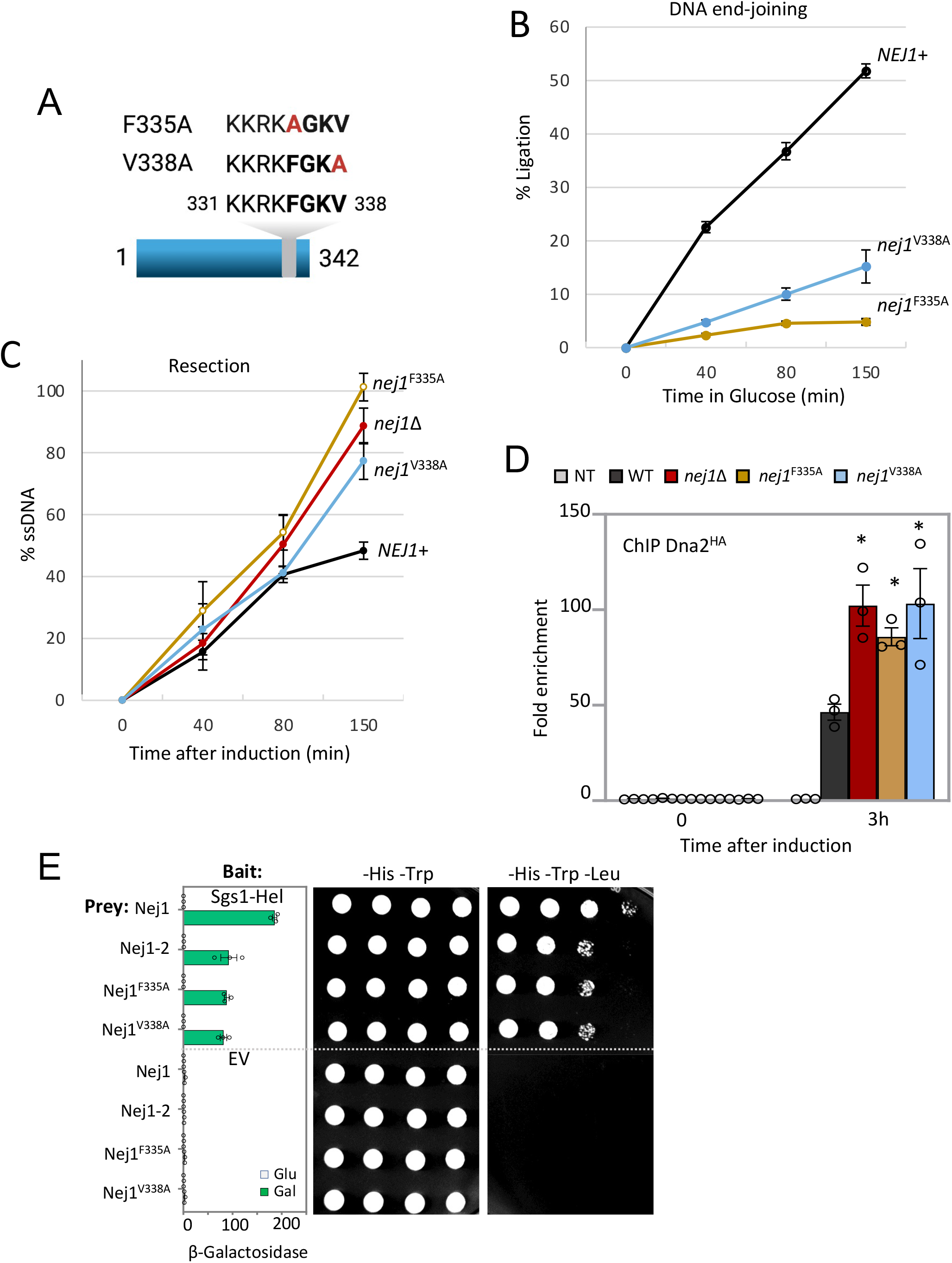
Nej1 Phenylalanine-335 and Valine-338 promote NHEJ by stimulating end-ligation and inhibiting Dna2. **(A)** Schematic representation of Nej1 C-terminal mutants *nej1*^F335A^ and *nej1*^V338A^ used in this study. **(B)** qPCR-based ligation assay of DNA at HO DSB, as measured by % Ligation, at 0, 40, 80 and 150 min in cycling cells in glucose post DSB. Strains used were wild type (JC-727), *nej1*^F335A^ (JC-2648) and *nej1*^V338A^ (JC-2659). **(C)** qPCR-based resection assay of DNA 0.15kb away from the HO DSB, as measured by % ssDNA, at 0, 40, 80 and 150 mins in cycling cells in wild type (JC-727), *nej1*Δ (JC-1342), *nej1*^F335A^ (JC-2648) and *nej1*^V338A^ (JC-2659). **(D)** Enrichment of Dna2^HA^ at 0.6kb from DSB 0 hour (no DSB induction) and 3 hours after DSB induction in wild type (JC-4117), *nej1*Δ (JC-4118), *nej1*^F335A^ (JC-5660), *nej1*^V338A^ (JC-5657) and no tag control (JC-727) was determined. The fold enrichment is normalized to recovery at the SMC2 locus. **(E)** Y2H analysis of Nej1 constructs fused to HA-AD; and Sgs1-Helicase core fused to lexA-DBD was performed in wild type cells (JC-1280) using a quantitative β-galactosidase assay and a drop assay on drop-out (-His, -Trp, -Leu) selective media plates. For all the experiments - error bars represent the standard error of three replicates. Significance was determined using 1-tailed, unpaired Student’s t test. All strains compared are marked (P<0.05*; P<0.01**; P<0.001***) unless specified with a line, strains are compared to WT.

We next measured 5’ resection and Dna2 recovery in the two mutants. Both *nej1*^F335A^ and *nej1*^V338A^ showed increased resection similarly to *nej1Δ* mutants, with *nej1*^F335A^ having the largest increase (Fig. 4C). Furthermore, this correlated with elevated levels of Dna2 recovered by ChIP at the DSB in both *nej1*^F335A^ and *nej1*^V338A^ mutants (Fig. 4D).

Previous work by our lab demonstrated that Nej1 regulated resection by inhibiting the binding of Dna2 to Sgs1 and Mre11. This prompted us to investigate whether these mutants showed altered binding to Mre11 or Sgs1 by Y2H. When expressed in prey vectors, Nej1-2 and both point mutants interacted with Mre11-C terminus similarly to wild type Nej1 (Fig. S2B-C). By contrast, the binding between Nej1 and Sgs1-helicase decreased significantly in every mutant (Fig. 4E and S2D). These results support a model wherein increased Dna2 at the DSB occurs because Nej1-Sgs1 binding decreases, and is consistent with earlier work showing Dna2-Sgs1 binding increased in *nej1*Δ mutants [3].

Of note, in addition to Ku, Mre11 is important for Nej1 recruitment [3]. Thus, the Y2H performed with Mre11-C and Nej1 complement ChIP data. We observed no disruption to interactions between these Nej1 mutant proteins and Mre11-C or in the recovery of Nej1 or other NHEJ factor at DSBs when *nej1*^F335A^ and *nej1*^V338A^ were expressed *in vivo* (Fig. S3A and [11]).

### 3.5 Nej1 Phenylalanine-335 is essential for end-ligation

As mentioned, although Ku and Nej1 interact, they appear to inhibit resection through different mechanisms. Ku protects DNA ends and inhibits resection mediated by Exo1, whereas increased resection in *nej1*Δ mutants depended on Dna2 but not Mre11 and Exo1 nuclease activities (Fig. 5A). The C-terminal region of Nej1 has overlapping functions as it both stimulates Lif1-Dnl4 and inhibit Dna2-mediated resection. Thus, inhibiting Dna2 activity should allow us to distinguish regions or residues in Nej1 intrinsically important for end-joining directly vs those that promote end-joining indirectly through down-regulating resection. Thus, we utilized nuclease dead *dna2*-1 (P504→S) as a tool to see whether we could restore end-joining in any of the *nej1* mutants. We first determined whether *dna2*-1 could reverse the hyper-resection of *nej1*Δ mutants. Cell harboring *nej1*Δ *dna2*-1 showed a complete defect in resection that was significantly below wild type (Fig 5B). By contrast, increased resection in *ku70*Δ mutant was not reversed in combination with *dna2*-1, but was Exo1 dependent (Fig 5B). This result is important as it demonstrates the unique antagonistic relationship of Nej1 and Dna2; and it also demonstrates that the impact of *dna2*-1 on resection is not dominant over all resection processes in general or resection mediated by other nucleases. Like in *nej1*Δ mutant, the increased resection in the C-terminal *nej1* mutants was reversed when *dna2*-1 was introduced (Fig. 3B, 4C, 5C-D).

**Figure 5.**
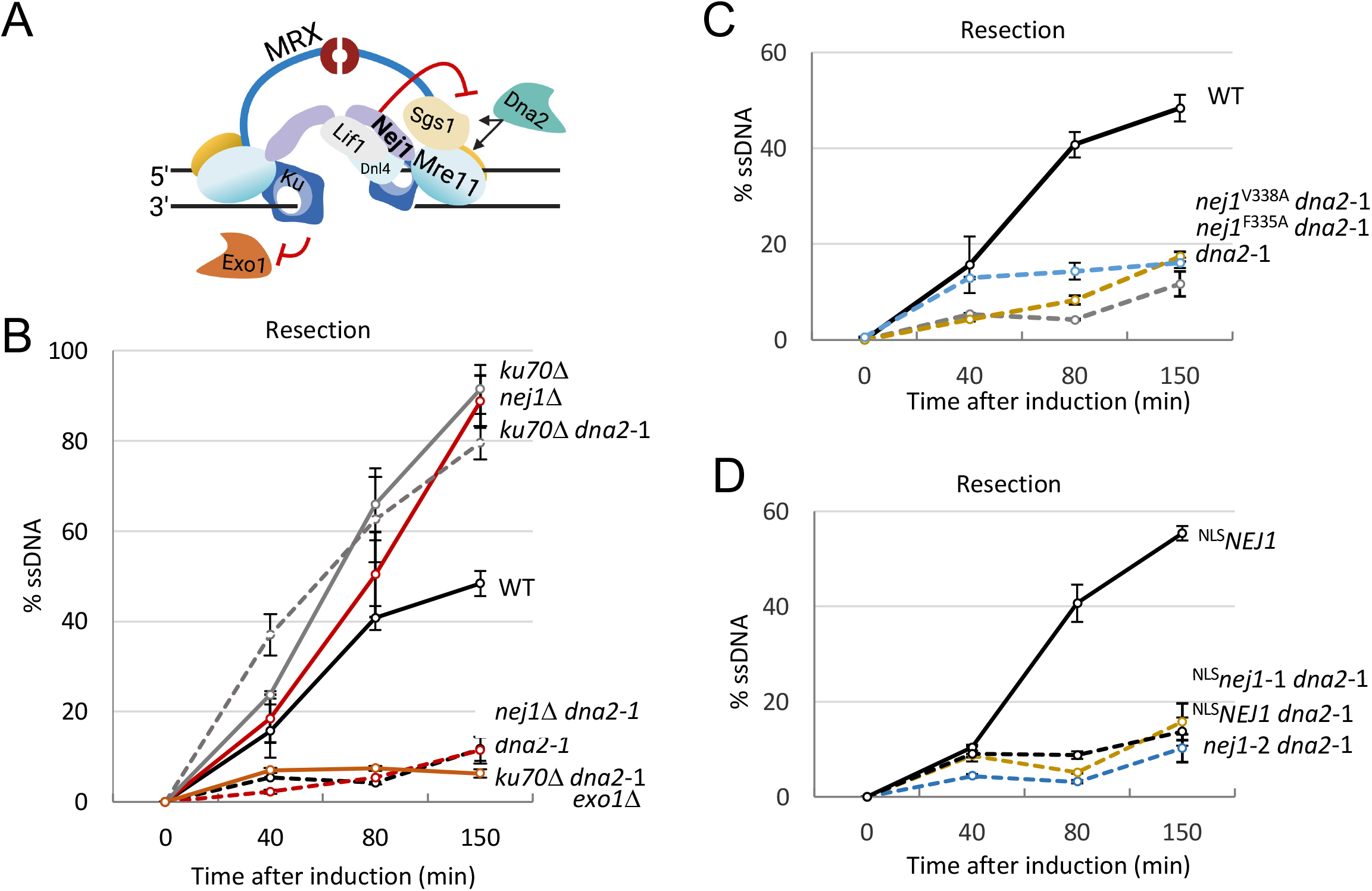
Interplay of Dna2 and Nej1 C-terminus at DSB. **(A)** Model depicting the antagonistic relationship of Ku-Exo1 and Nej1-Dna2 at DSB. **(B - D)** qPCR-based resection assay of DNA 0.15kb away from the HO DSB, as measured by % ssDNA, at 0, 40, 80 and 150 min post DSB induction in cycling cells in wild type (JC-727), *nej1*Δ (JC-1342), *dna2*-1 (JC-5655), *nej1*Δ *dna2*-1 (JC-5670), *ku70*Δ (JC-1904), *ku70*Δ *dna2*-1 (JC-5942), *ku70*Δ *dna2*-1 *exo1*Δ (JC-6025), *nej1*^F335A^ *dna2*-1 (JC-5804), *nej1*^V338A^ *dna2*-1 (JC-5806), *nej1*-1 *dna2*-1 (JC-5874) and *nej1*-2 *dna2*-1 (JC-5870). For all the experiments - error bars represent the standard error of three replicates.

As resection decreased to below wild type when the *nej1* mutants were combined with *dna2*-1, we were encouraged that *dna2*-1 would be an excellent genetic tool to distinguish direct from indirect Nej1 functions in NHEJ. Indeed, the rate of end-joining in *dna2*-1 mutant cells increased above wild type and was dependent on Ku, Nej1 and the other core NHEJ factors (Fig. 6A and S3B). The end-joining defect in *nej1*^V338A^ mutant cells was reversed in combination with *dna2*-1 (Fig. 4B and 6B), however in *nej1*^F335A^ *dna2*-1 end-joining was as defective as in *nej1*^F335A^ mutants. Furthermore, the end-joining defect in ^NLS^*nej1*-1 was also reversed in combination with *dna2*-1 (Fig. 1F and 6C). These data show that if resection is inhibited in ^NLS^*nej1*-1, then end-joining can proceed. Thus, the binding of Nej1 to ssDNA and dsDNA is not intrinsically necessary for stimulating ligation.

**Figure 6.**
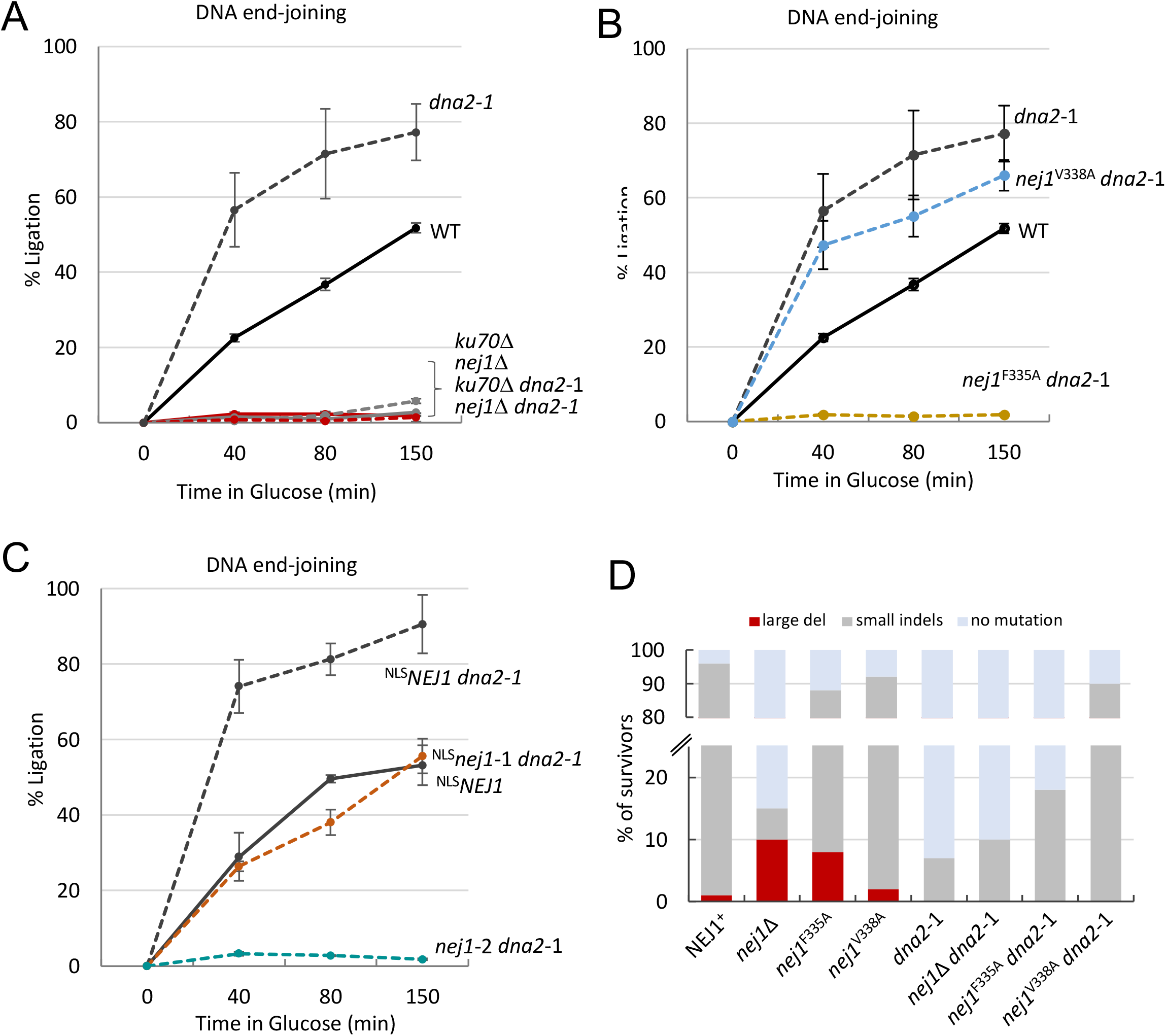
Nej1 Phenylalanine-335 is essential for end-ligation. **(A - C)** qPCR-based ligation assay of DNA at HO DSB, as measured by % Ligation, at 0, 40, 80 and 150 min in cycling cells in glucose post DSB. Strains used were wild type (JC-727), *nej1*Δ (JC-1342), *dna2*-1 (JC-5655), *nej1*Δ *dna2*-1 (JC-5670), *ku70*Δ (JC-1904), *ku70*Δ *dna2*-1 (JC-5942), *nej1*^F335A^ *dna2*-1 (JC-5804), *nej1*^V338A^ *dna2*-1 (JC-5806), *NEJ1*^*+*^ (JC-3193), *NEJ1*^*+*^ *dna2*-1 (JC-5871), *nej1*-1 *dna2*-1 (JC-5874) and *nej1*-2 *dna2*-1 (JC-5870). **(D)** Percentage cell survival upon chronic HO induction in wild type (JC-727), *nej1*Δ (JC-1342), *dna2*-1 (JC-5655), *nej1*Δ *dna2*-1 (JC-5670), *nej1*^F335A^ (JC-2648), *nej1*^F335A^ *dna2*-1 (JC-5804), *nej1*^V338A^ (JC-2659) and *nej1*^V338A^ *dna2*-1 (JC-5806). For all the experiments - error bars represent the standard error of three replicates.

Similar to *nej1*-2, large deletion developed in *nej1*^F335A^, and to a lesser degree in *nej1*^V338A^ survivors (Fig. 6D). Survival increased in *dna2*-1 mutants above wild type, which is consistent with having increased end-joining repair occur as a consequence of decreased resection (Fig. 6A). Moreover, large deletions were not observed in any *nej1 dna2*-1 double mutant combinations (Fig. 6D). While end-joining was restored in ^NLS^*nej1*-1 *dna2*-1 and *nej1*^V338A^ dna2-1, large deletions did not develop at the break because resection was also reduced (Fig. 6D). These data strongly suggest that the loss of resection inhibition, rather than the loss of end-joining stimulation, drives the formation of large deletions in *nej1* mutants.

## 4. DISCUSSION

### 4.1 Nej1 promotes NHEJ through overlapping C-terminal functions

Nej1 has multiple properties that impact repair pathway choice. Characterizing *nej1* mutants allowed us to determine how specific regions between K331 and V338 of Nej1 C-terminus function to promote NHEJ (Fig. 7). The C-terminus of Nej1 binds DNA, inhibits resection, and drives DNA end-joining, but is dispensable for end-bridging. Nej1 ChIP in ^NLS^*nej1*ΔC (1-330 aa) supported our earlier work showing that the Lif1 interacting region, FGKV (335-338), was not important for Nej1 recruitment to the DSB [21]. Further, the data shown here indicate that the KKRK (331-334) region was also dispensable for Nej1 recruitment. Nej1 recovery at the DSB in ^NLS^*nej1*-1 supported this too, which was interesting given KKRK is the signal localizing Nej1 to the nucleus and for interacting with ssDNA and dsDNA (Fig. 1C; [21, 25], while FGKV (335-338) region was expendable for DNA binding (Fig. 2C-D). Taken together, after Nej1 recruitment, the ability of KKRK to bind ssDNA could promote end-joining by stabilizing DNA ends, as HO-DSBs have compatible ssDNA overhangs used in direct ligation by NHEJ [31].

**Figure 7.**
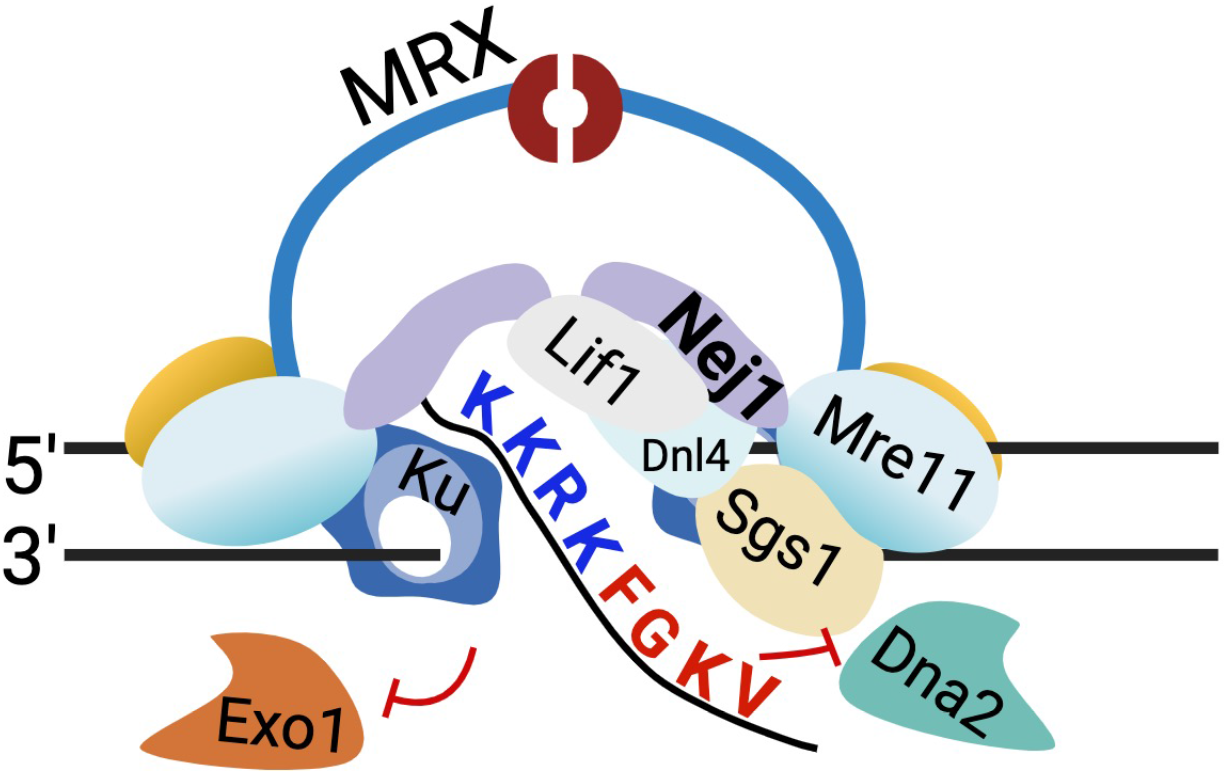
Model depicting the role of Nej1 C-terminus at DSB. NHEJ complex end-ligates, where Nej1 stimulates end-ligation via its C-terminal region. The ‘KKRK’ motif binds ssDNA overhangs and facilitates ligation and the ‘FGKV’ motif directly stimulates ligation by physically interacting with Lif1-Dnl4 and inhibiting Dna2 at DSB. The model depicts the antagonistic relationship of end-joining complex and Dna2 nuclease.

Double mutant combinations of the NHEJ factors deleted in combination with *dna2*-1 showed that an increase in the frequency of large deletions correlated with increased resection rather than decreased end-joining [11]. End-joining was abrogated when any one of the core NHEJ factors was deleted, however *ku70*Δ *dna2*-1 was the only *dna2*-1 double mutant combination that produced survivors with large deletions (Table 1). As mentioned, resection in *ku70*Δ was not Dna2 dependent (Fig. 5B). By contrast, increased resection in *nej1*Δ was Dna2 dependent, and *nej1*Δ *dna2*-1 double mutants showed decreased resection and no large deletions. This trend held when comparing ^NLS^*nej1*-1 and *nej1*-2. Both resection and the frequency of deletions increased in *nej1*-2 compared to ^NLS^*nej1*-1 mutants (Fig. 3B and Table 1), indicating that FGKV, but not KKRK, was important for inhibiting Dna2-mediated resection (Fig. 3C).

### 4.2 Nej1 promotes end-joining directly and indirectly

Comparing repair events in F335A and V338A revealed that the frequency of genomic deletion also depended on intrinsic end-joining potential. Both *nej1*^F335A^ and *nej1*^V338A^ showed increased resection and decreased end-joining (Fig. 4B-C), however *nej1*^F335A^ mutants showed a higher frequency of large deletions compared to *nej1*^V338A^. While both F335 and V338 were important for Dna2 inhibition (Fig. 4D), when resection was reduced by *dna2*-1, only *nej1*^V338A^ *dna2*-1 double mutants demonstrated increased end-joining and increased survival (Fig. 6B and D), *nej1*^F335^ *dna2* -1 did not. Thus, properties within F335, unrelated to inhibiting resection, were intrinsic important in end-joining repair. This C-terminal region shows sequence conservation from Nej1 in yeast to XLF in humans, as the both the KKRK motif and the invariant phenylalanine (F335 in Nej1) are present in almost all organisms [21-25, 29]. These observations also help clarify perplexing results wherein *nej1*^V338A^ mutants showed proficient Dnl4 -ligase dependent repair in plasmid rejoining assays, but a marked reduction in viability after HO-DSB induction (Table 1, [21]). Perhaps a DSB within a plasmid is less accessible to nucleases and resection compared to a DSB located at the HO cut site in the genome. The *nej1*^V338A^ mutant is deficient in inhibiting resection, but proficient in end-joining, and exhibits hallmark separation-of-function qualities. Reduced viability *nej1*^V338A^ single mutants stemmed from indirect defects in end-joining attributed to increased resection. By investigating the mating-type of *nej1*^V338A^ survivors, we observed that the majority of repair events proceeded by Dnl4-dependent error-prone direct ligation, similarly to events in *NEJ1*+ (Fig. 6D and S3B, Table1).

### 4.3 Nej1 and Dna2, a tug of war between end-joining or end-resection

The antagonistic relation of Nej1 and Dna2 regulates DSB repair pathway choice. Nej1 is a core NHEJ factor that stimulates end-joining and inhibits Dna2-mediated 5’ resection. Whereas Dna2 is a central resection factor that inhibit end-joining repair. The inhibition of end-joining by Dna2 was distinctive, as the frequency of end-joining did not increase by disruptions in other resection promoting factors (Fig. S3C). Consistent with this model, the recovery of Ku70 and Nej1 by ChIP also increased at DSBs in *dna2*-1 mutants [13].

Indeed, survival of *dna2*-1 mutants increased in engineered cells where end-joining was the only pathway available to repair the HO-DSB. However, the core NHEJ factors remained essential as survival rates as observed by comparing *nej1*Δ and *nej1*Δ *dna2*-1, although large deletions did not develop in double mutant survivors because resection was also compromised (Fig. S3D). Similar trends were observed when *dna2*-1 was combined with deletions in the other core NHEJ factors. All NHEJ mutant combinations showed reduced end-joining repair regardless of resection level. This demonstrates the essentiality of all the core NHEJ factors, which cannot be surmounted by alterations in resection. These insights from yeast point to reasons underlying human XLF mutations in genome instability and immunodeficient diseases [33].

## Acknowledgements

This work was supported by operating grants from CIHR MOP-82736; MOP-137062 and NSERC 418122 awarded to J.A.C. We acknowledge the resources provided by the Live Cell Imaging Laboratory. The Nikon Ti Eclipse inverted epi-fluorescence microscope system was purchased with funds from the International Microbiome Centre, which is supported by the Cumming School of Medicine at University of Calgary, Western Economic Diversification (WED) and Alberta Economic Development and Trade (AEDT), Canada. We thank Dr. Gareth Williams for kindly providing us the DNA substrates and FP measuring machine PHERAstar to perform our DNA binding assays.

## Supplemental data

**Supplementary Figure S1.**
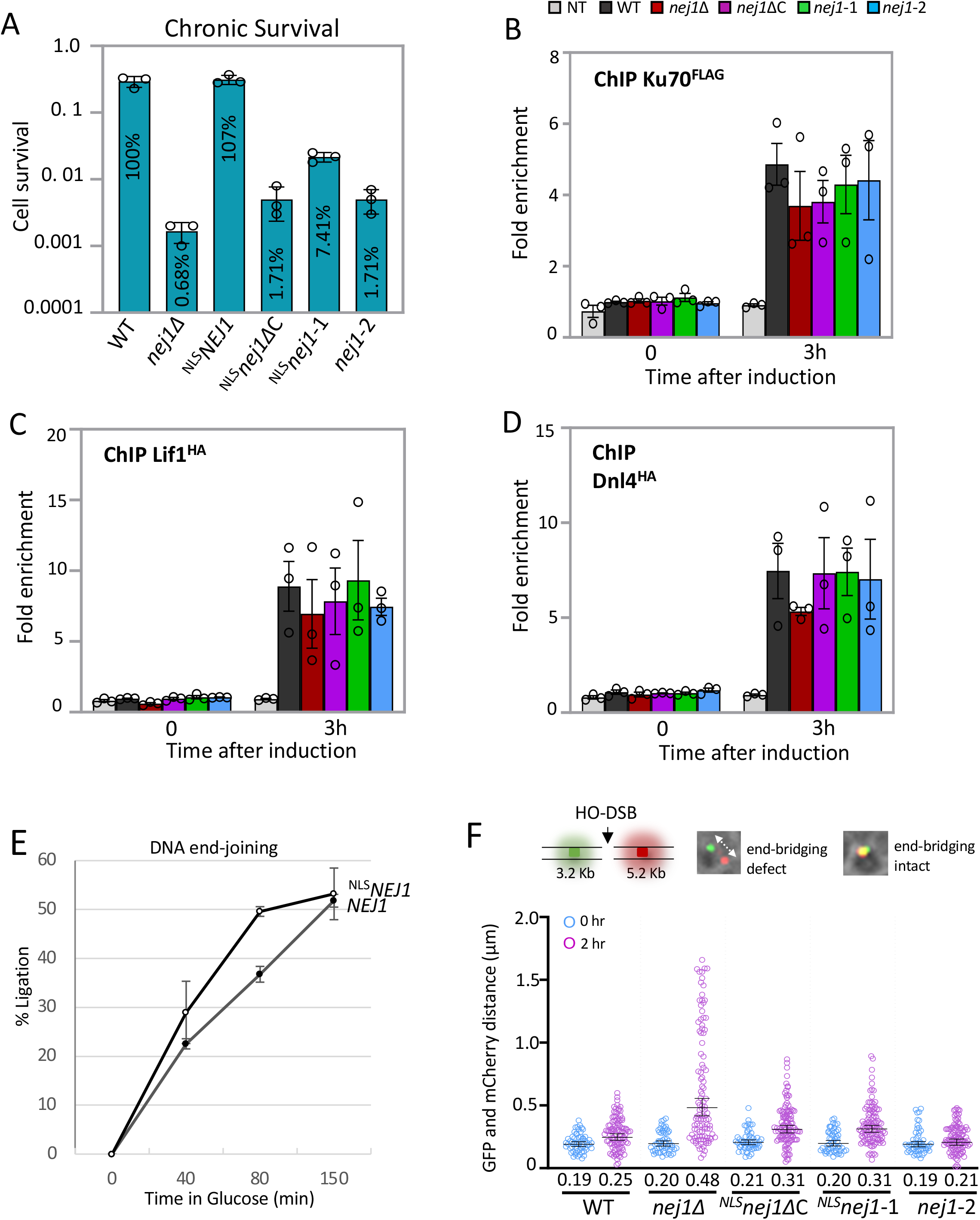
**(A)** Percentage cell survival upon chronic HO induction in wild type (JC-727), *nej1*Δ (JC-1342), *NEJ1*^*+*^ (JC-3193), *nej1*ΔC (JC-3194), *nej1*-1 (JC-3215) and *nej1*-2 (JC-3134). **(B, C and D)** Enrichment of Ku70^Flag^ / Lif1^HA^ / Dnl4^HA^ at DSB, at 0 and 3 hours, in wild type (JC-3964 / 3319 / 5672), *nej1*Δ (JC-5941 / 3346 / 5648), *nej1*ΔC (JC-5926 / 5852 / 5860), *nej1*-1 (JC-5928 / 5855 / 5864), *nej1*-2 (JC-5930 / 5853 / 5863) and no tag control (JC-727). **(E)** qPCR-based ligation assay of DNA at HO DSB, as measured by % Ligation, at 0, 40, 80 and 150 min in cycling cells in glucose post DSB. Strains used were wild type (JC-727) and *NEJ1*^*+*^ (JC-3193). **(F)** Representative image of yeast cells with tethered (co-localized GFP and mCherry) and untethered (distant GFP and mCherry) ends. Scatter plot showing the End-bridging of DSB ends, at 0 and 2 hours, as measured by the distance between the GFP and mCherry foci in wild type (JC-4066), *nej1*Δ (JC-4364), *nej1*ΔC (JC-5875), *nej1*-1 (JC-5876) and *nej1*-2 (JC-5877). The Geometric mean (GM) distance for each sample is specified under the respective sample data plot. Significance was determined using Kruskal-Wallis and Dunn’s multiple comparison test. For all the experiments - error bars represent the standard error of three replicates.

**Supplementary Figure S2.**
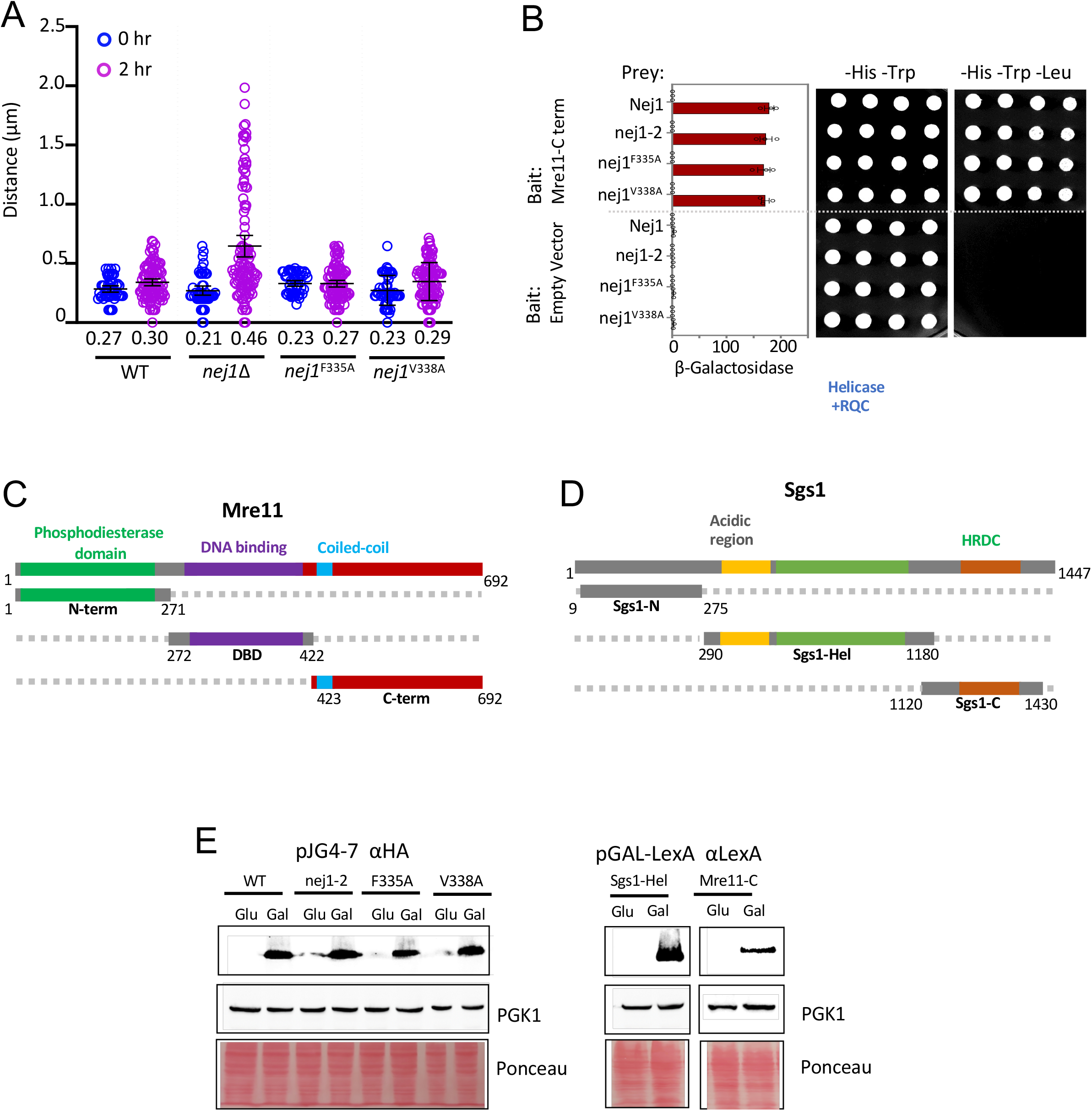
**(A)** Scatter plot showing the End-bridging of DSB ends, at 0 and 2 hours, as measured by the distance between the GFP and mCherry foci in wild type (JC-4066), *nej1*Δ (JC-4364), *nej1*^F335A^ (JC-4073), and *nej1*^V338A^ (JC-4074). The Geometric mean (GM) distance for each sample is specified under the respective sample data plot. Significance was determined using Kruskal-Wallis and Dunn’s multiple comparison test. **(B)** Y2H analysis of Nej1 constructs fused to HA-AD; and Mre11-C terminal region fused to lexA-DBD was performed in wild type cells (JC-1280) using a quantitative β-galactosidase assay and a drop assay on drop-out (-His, -Trp, -Leu) selective media plates. **(C)** Schematic representation of Mre11 and its functional domains. In green is the N-terminus (1-271 aa) of Mre11 that contains the phosphodiesterase domain, in purple is the DNA binding domain (272–422 aa) and in dark red is the C-terminal region (423–692 aa) containing the Rad50 binding site. This is adapted from Mojumdar et al., 2019 [3]. **(D)** Schematic representation of Sgs1 and its functional domains. The N-terminal region (9-275 aa) is in gray, the helicase core (290-1180 aa) is in green, and the C-terminal helicase and RNaseD C-terminal (HRDC) domain (1120-1430 aa) is in orange. This is adapted from Mojumdar et al., 2019 [3]. **(E)** Expression of proteins upon galactose induction. Western blots showing the level of expression of proteins fused to LexA and HA tag upon galactose induction in WT. PGK1 was stained as a loading control. For all the experiments - error bars represent the standard error of three replicates.

**Supplementary Figure S3.**
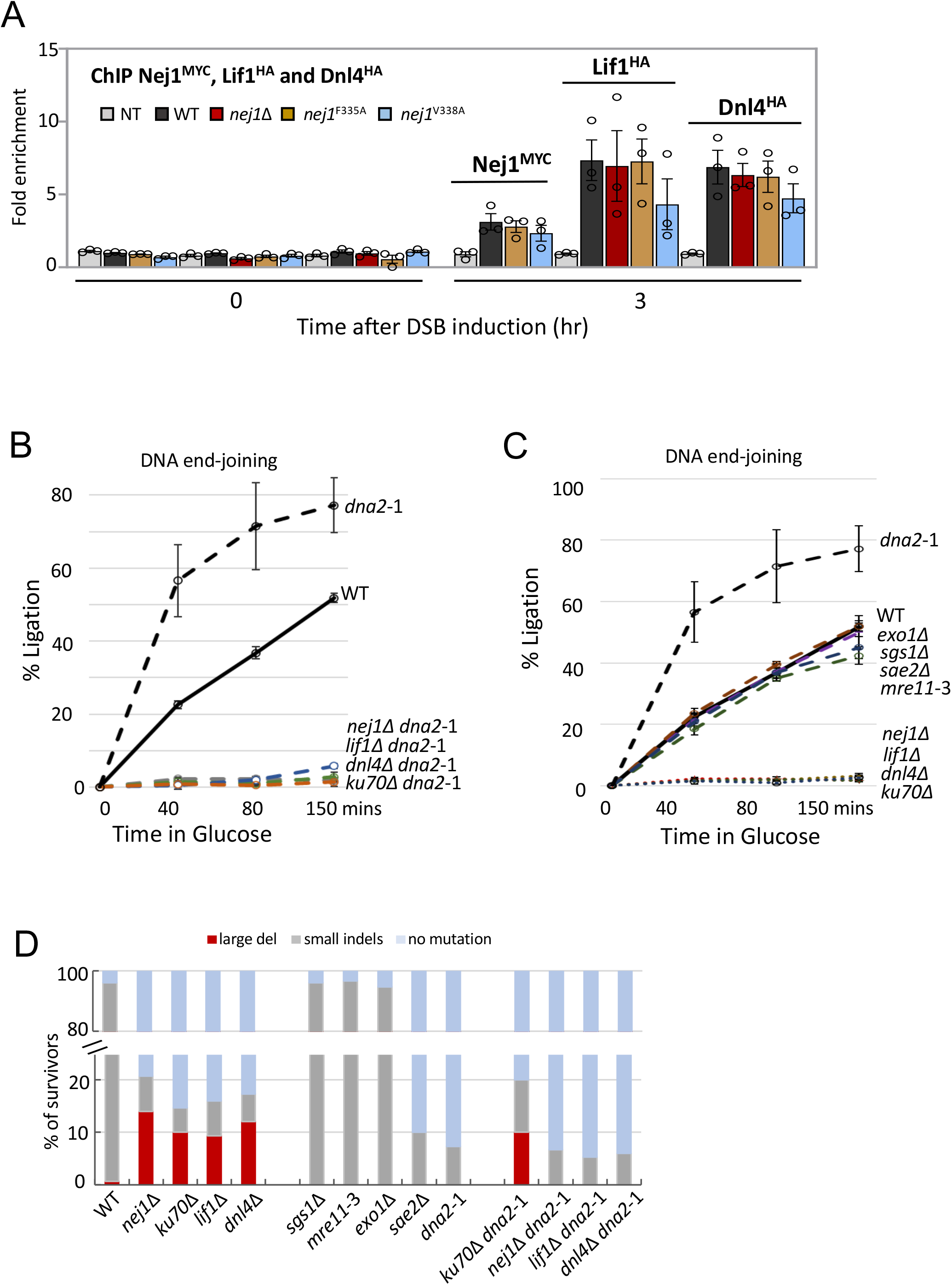
**(A)** Enrichment of Nej1^MYC^ / Lif1^HA^ / Dnl4^HA^ at DSB, at 0 and 3 hours, in wild type (JC-1687 / 3319 / 5672), *nej1*Δ (JC- / 3346 / 5648), *nej1*^F335A^ (JC-3133 / 3816 / 5649), *nej1*^V338A^ (JC-3160 / 3828 / 5650) and no tag control (JC-727). **(B)** qPCR-based ligation assay of DNA at HO DSB, as measured by % Ligation, at 0, 40, 80 and 150 min in cycling cells in glucose post DSB. Strains used were wild type (JC-727), *dna2*-1 (JC-5655), *lif1*Δ (JC-1343), *lif1*Δ *dna2*-1 (JC-5890), *dnl4*Δ (JC-3290) and *dnl4*Δ *dna2*-1 (JC-5800). **(C)** qPCR-based resection assay of DNA 0.15kb away from the HO DSB, as measured by % ssDNA, at 0, 40, 80 and 150 mins in cycling cells in wild type (JC-727), *nej1*Δ (JC-1342), *lif1*Δ (JC-1343), *ku70*Δ (JC-1904), *dnl4*Δ (JC-3290), *sgs1*Δ (JC-3757), *exo1*Δ (JC-3767), *mre11*-3 (JC-5372), *sae2*Δ (JC-5673) and *dna2*-1 (JC-5655). **(D)** Percentage cell survival upon chronic HO induction in wild type wild type (JC-727), *nej1*Δ (JC-1342), *lif1*Δ (JC-1343), *ku70*Δ (JC-1904), *dnl4*Δ (JC-3290), *sgs1*Δ (JC-3757), *exo1*Δ (JC-3767), *mre11*-3 (JC-5372), *sae2*Δ (JC-5673) and *dna2*-1 (JC-5655), *ku70*Δ *dna2*-1 (JC-5942), *nej1*Δ *dna2*-1 (JC-5670), *lif1*Δ *dna2*-1 (JC-5890), and *dnl4*Δ *dna2*-1 (JC-5800). For all the experiments - error bars represent the standard error of three replicates.

**Table S1 Yeast Strains:**
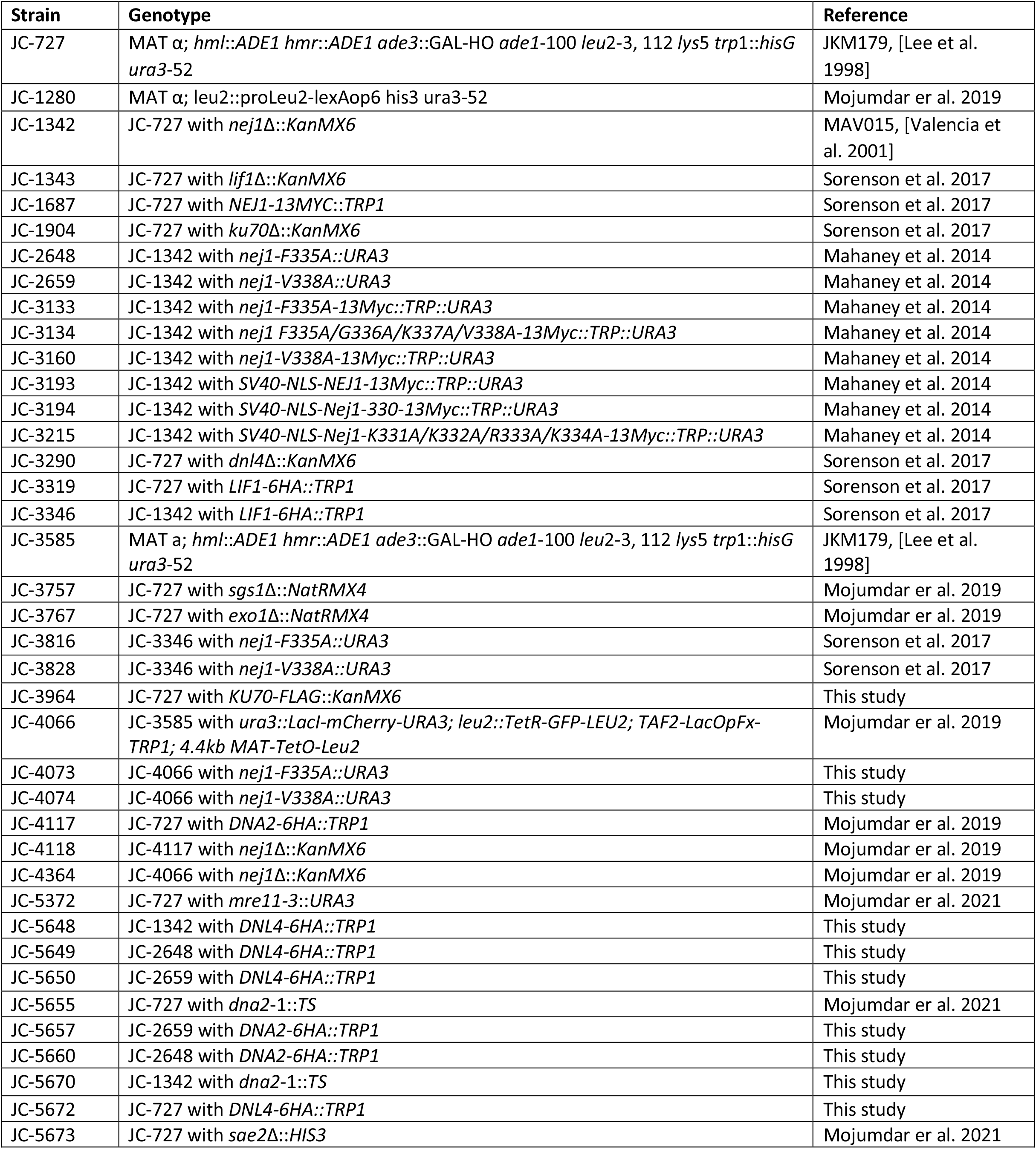
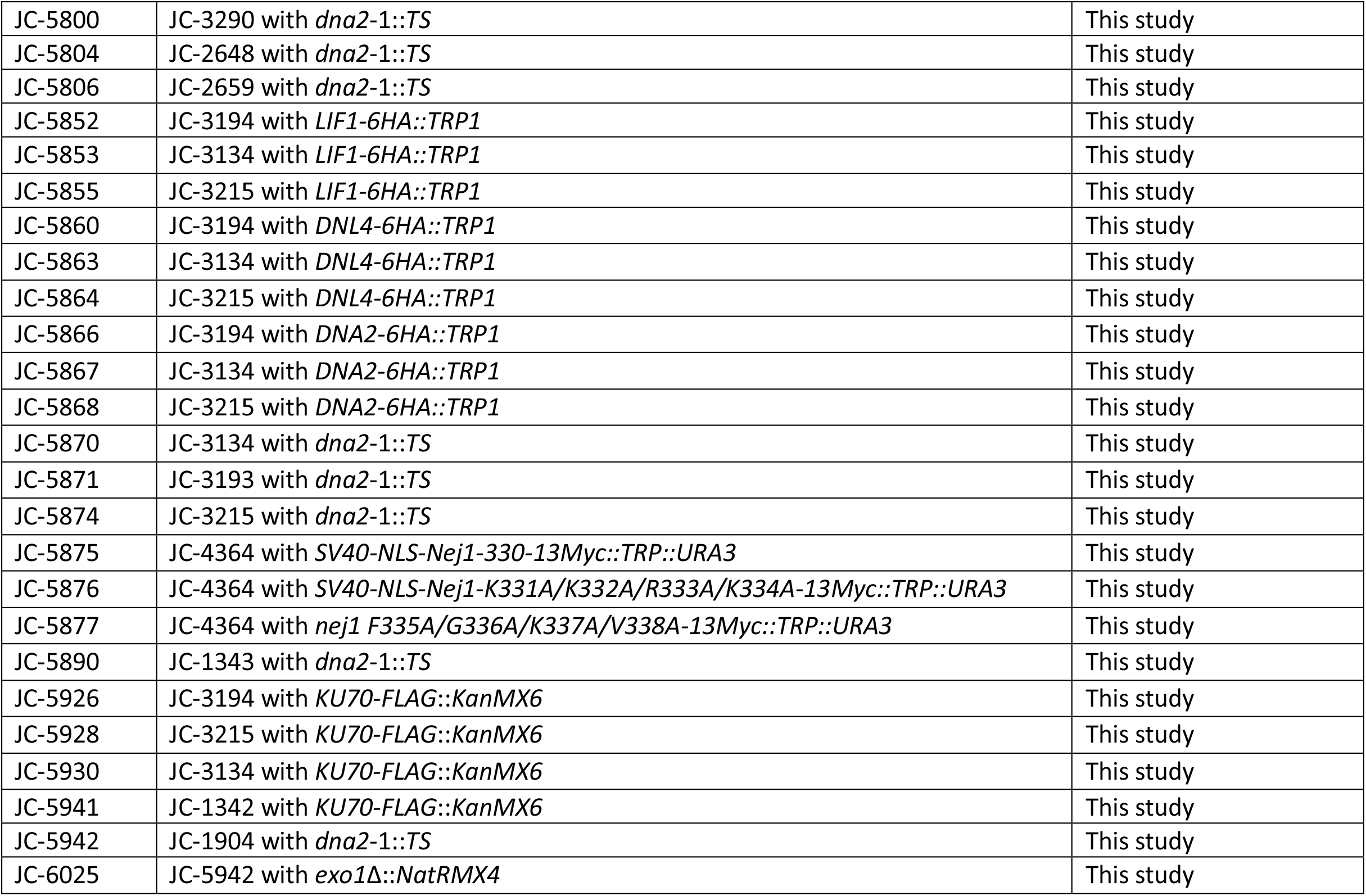
The yeast strains used in this study are outlined.

**Table S2 Plasmids:** Plasmids used in this study are listed below.

| Plasmid number | Description |
| --- | --- |
| J-965 | pGAL-LexA |
| J-1493 | pJG4-6 |
| J-123 | J-1493 with Nej1 |
| J-127 | J-1493 with Nej1 (F335A) |
| J-130 | J-1493 with Nej1 (V338A) |
| J-136 | J-1493 with Nej1 (FGKV-4A) |
| J-1044 | J-965 with Sgs1-(290-1180) Helicase |
| J-1872 | J-965 with Mre11-(423-692) C-term |
| J-1917 | J-1493 with Nej1 (1-330) |
| J-1918 | J-1493 with Nej1 (KKRK-4A) |

**Table S3 Primers:**
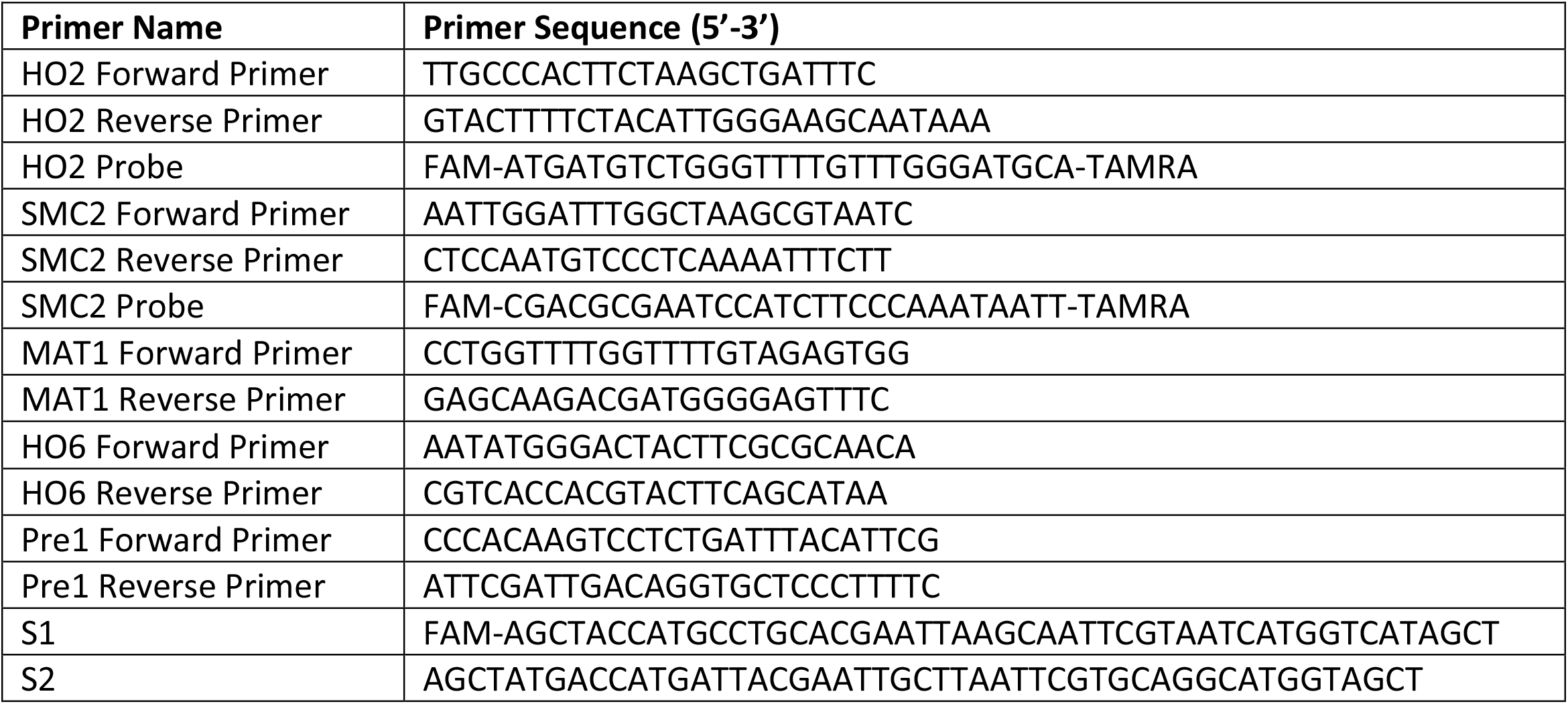
Table of Primers and Probes used in these studies.

**Table S4:**
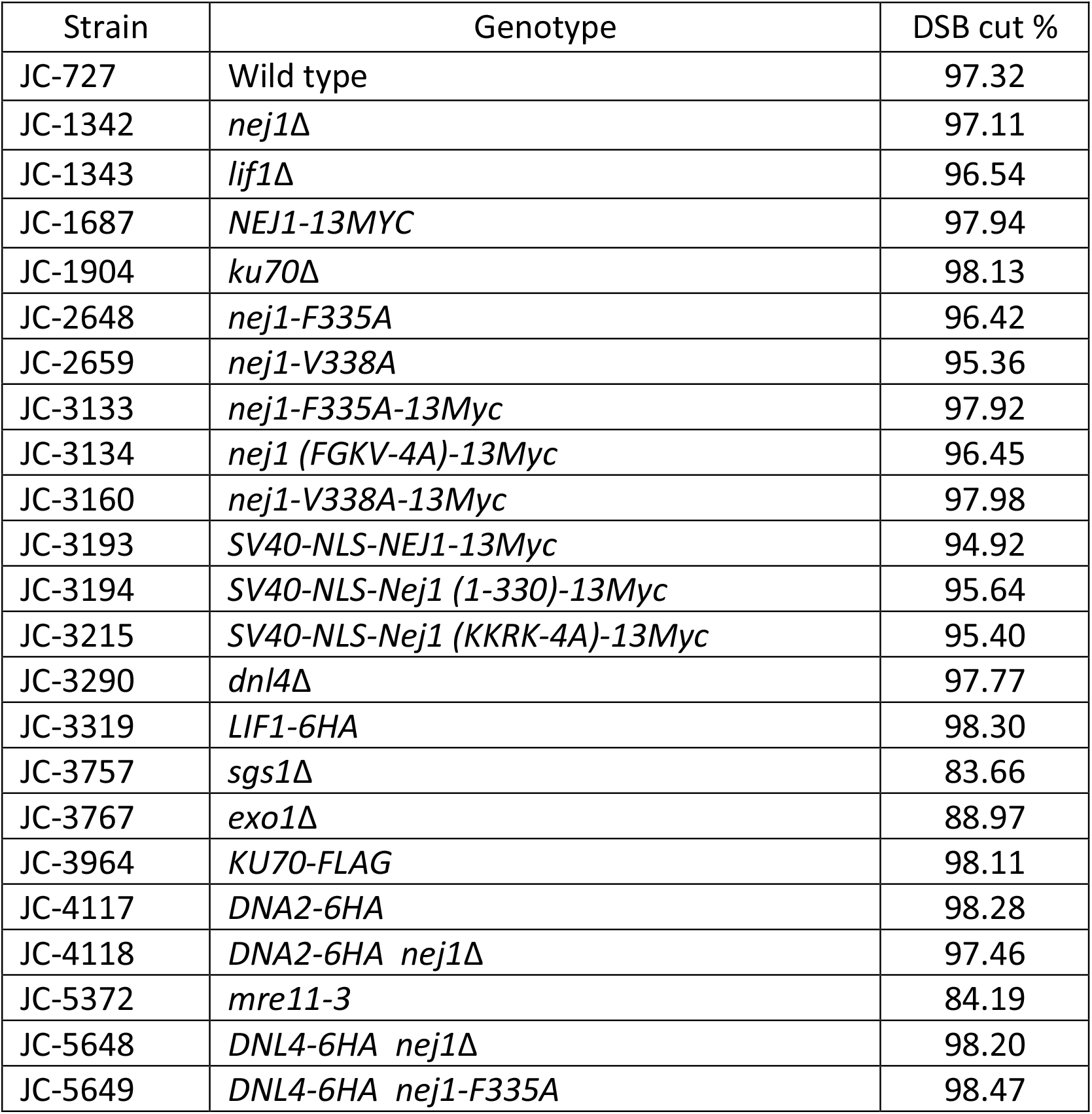
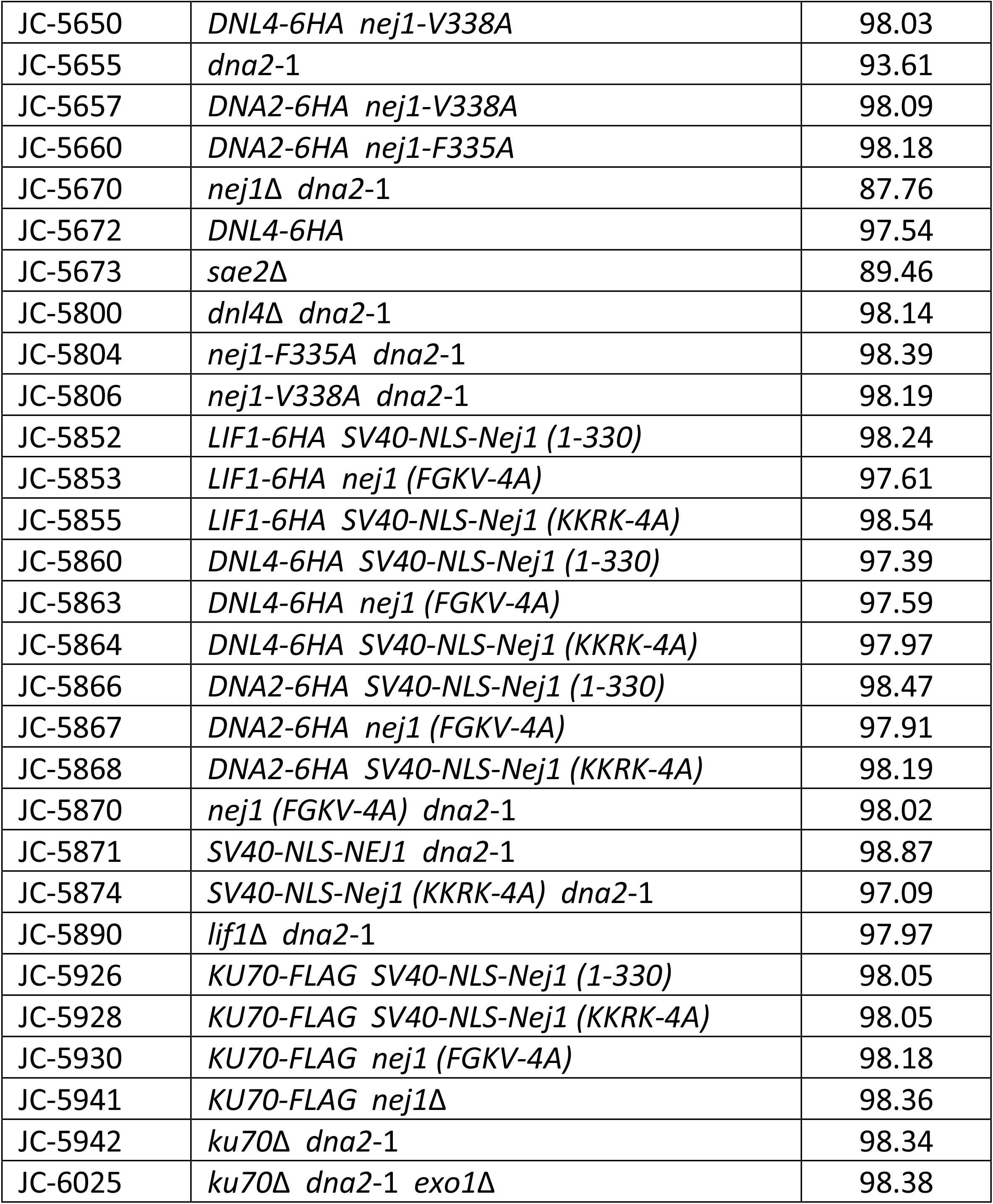
Table of DSB cut efficiency.

| Strain | Genotype | DSB cut % |
| --- | --- | --- |
| JC-727 | Wild type | 97.32 |
| JC-1342 | *nej1*Δ | 97.11 |
| JC-1343 | *lif1*Δ | 96.54 |
| JC-1687 | *NEJ1-13Myc* | 97.94 |
| JC-1904 | *ku70*Δ | 98.13 |
| JC-2648 | *nej1-F335A* | 96.42 |
| JC-2659 | *nej1-V338A* | 95.36 |
| JC-3133 | *nej1-F335A-13Myc* | 97.92 |
| JC-3134 | *nej1 (FGKV-4A)-13Myc* | 96.45 |
| JC-3160 | *nej1-V338A-13Myc* | 97.98 |
| JC-3193 | *SV40-NLS-NEJ1-13Myc* | 94.92 |
| JC-3194 | *SV40-NLS-Nej1 (1-330)-13Myc* | 95.64 |
| JC-3215 | *SV40-NLS-Nej1 (KKRK-4A)-13Myc* | 95.40 |
| JC-3290 | *dnl4*Δ | 97.77 |
| JC-3319 | *LIF1-6HA* | 98.30 |
| JC-3757 | *sgs1*Δ | 83.66 |
| JC-3767 | *exo1*Δ | 88.97 |
| JC-3964 | *KU70-FLAG* | 98.11 |
| JC-4117 | *DNA2-6HA* | 98.28 |
| JC-4118 | *DNA2-6HA nej1*Δ | 97.46 |
| JC-5372 | *mre11-3* | 84.19 |
| JC-5648 | *DNL4-6HA nej1*Δ | 98.20 |
| JC-5649 | *DNL4-6HA nej1-F335A* | 98.47 |
| JC-5650 | *DNL4-6HA nej1-V338A* | 98.03 |
| JC-5655 | *dna2-1* | 93.61 |
| JC-5657 | *DNA2-6HA nej1-V338A* | 98.09 |
| JC-5660 | *DNA2-6HA nej1-F335A* | 98.18 |
| JC-5670 | *nej1Δ dna2-1* | 87.76 |
| JC-5672 | *DNL4-6HA* | 97.54 |
| JC-5673 | *sae2Δ* | 89.46 |
| JC-5800 | *dnl4Δ dna2-1* | 98.14 |
| JC-5804 | *nej1-F335A dna2-1* | 98.39 |
| JC-5806 | *nej1-V338A dna2-1* | 98.19 |
| JC-5852 | *LIF1-6HA SV40-NLS-Nej1 (1-330)* | 98.24 |
| JC-5853 | *LIF1-6HA nej1 (FGKV-4A)* | 97.61 |
| JC-5855 | *LIF1-6HA SV40-NLS-Nej1 (KKRK-4A)* | 98.54 |
| JC-5860 | *DNL4-6HA SV40-NLS-Nej1 (1-330)* | 97.39 |
| JC-5863 | *DNL4-6HA nej1 (FGKV-4A)* | 97.59 |
| JC-5864 | *DNL4-6HA SV40-NLS-Nej1 (KKRK-4A)* | 97.97 |
| JC-5866 | *DNA2-6HA SV40-NLS-Nej1 (1-330)* | 98.47 |
| JC-5867 | *DNA2-6HA nej1 (FGKV-4A)* | 97.91 |
| JC-5868 | *DNA2-6HA SV40-NLS-Nej1 (KKRK-4A)* | 98.19 |
| JC-5870 | *nej1 (FGKV-4A) dna2-1* | 98.02 |
| JC-5871 | *SV40-NLS-NEJ1 dna2-1* | 98.87 |
| JC-5874 | *SV40-NLS-Nej1 (KKRK-4A) dna2-1* | 97.09 |
| JC-5890 | *lif1Δ dna2-1* | 97.97 |
| JC-5926 | *KU70-FLAG SV40-NLS-Nej1 (1-330)* | 98.05 |
| JC-5928 | *KU70-FLAG SV40-NLS-Nej1 (KKRK-4A)* | 98.05 |
| JC-5930 | *KU70-FLAG nej1 (FGKV-4A)* | 98.18 |
| JC-5941 | *KU70-FLAG nej1Δ* | 98.36 |
| JC-5942 | *ku70Δ dna2-1* | 98.34 |
| JC-6025 | *ku70Δ dna2-1 exo1Δ* | 98.38 |

## REFERENCES

1. L. Krejci, V. Altmannova, M. Spirek, X. Zhao, Homologous recombination and its regulation. Nucleic Acids Res. 40(13) (2012) 5795–5818. doi:10.1093/nar/gks270.

2. S. K. Radhakrishnan, N. Jette, S. P. Lees-Miller, Non-homologous end joining: emerging themes and unanswered questions. DNA Repair Amst. 17 (2014) 2–8.

3. A. Mojumdar, K. Sorenson, M. Hohl, M. Toulouze, S. P. Lees-Miller, K. Dubrana, J. H. J. Petrini, J. A. Cobb, Nej1 Interacts with Mre11 to Regulate Tethering and Dna2 Binding at DNA Double-Strand Breaks. Cell Rep. 28(6) (2019) 1564–1573. doi: 10.1016/j.celrep.2019.07.018.

4. P. L. Palmbos, D. Wu, J. M. Daley, T. E. Wilson, Recruitment of Saccharomyces cerevisiae Dnl4-Lif1 complex to a double-strand break requires interactions with Yku80 and the Xrs2 FHA domain. Genetics. 180 (2008) 1809–1819.

5. S. H. Teo, S.P. Jackson, Identification of Saccharomyces cerevisiae DNA ligase IV: involvement in DNA double-strand break repair. EMBO J. 16 (1997) 4788–4795.

6. T. E. Wilson, U. Grawunder, M. R. Lieber. Yeast DNA ligase IV mediates non-homologous DNA end joining. Nature. 388 (1997) 495–498.

7. G. Herrmann, T. Lindahl, P. Schar, Saccharomyces cerevisiae LIF1: a function involved in DNA double-strand break repair related to mammalian XRCC4. EMBO J. 17 (1998) 4188– 4198.

8. X. Chen, A. E. Tomkinson, Yeast Nej1 is a key participant in the initial end binding and final ligation steps of nonhomologous end joining. J. Biol. Chem. 286 (2011) 4931–4940.

9. M. Frank-Vaillant, S. Marcand, NHEJ regulation by mating type is exercised through a novel protein, Lif2p, essential to the ligase IV pathway. Genes Dev. 15(22) (2001) 3005–3012. doi: 10.1101/gad.206801.

10. M. Valencia, M. Bentele, M. B. Vaze, G. Herrmann, E. Kraus, S. E. Lee, P. Schär, J. E. Haber, NEJ1 controls non-homologous end joining in Saccharomyces cerevisiae. Nature. 414(6864) (2001) 666–669. doi: 10.1038/414666a.

11. K. S. Sorenson, B. L. Mahaney, S. P. Lees-Miller, J. A. Cobb, The non-homologous end-joining factor Nej1 inhibits resection mediated by Dna2-Sgs1 nuclease-helicase at DNA double strand breaks. J Biol Chem. 292(35) (2017) 14576–14586.

12. E. P. Mimitou, L. S. Symington, Ku prevents Exo1 and Sgs1-dependent resection of DNA ends in the absence of a functional MRX complex or Sae2. EMBO J. 29(19) (2010) 3358–3369.

13. A. Mojumdar, N. Adam, J. A. Cobb. Sgs1^BLM^independent role of Dna2^DNA2^ nuclease at DNA double strand break is inhibited by Nej1^XLF^. bioRxiv 2021.04.10.439283. doi: https://doi.org/10.1101/2021.04.10.439283D.

14. S. H. Bae, K. H. Bae, J. A. Kim, Y. S. Seo, RPA governs endonuclease switching during processing of Okazaki fragments in eukaryotes. Nature. 412(6845) (2001) 456–461. doi: 10.1038/35086609.

15. Z. Zhu, W. H. Chung, E. Y. Shim, S. E. Lee, G. Ira, Sgs1 helicase and two nucleases Dna2 and Exo1 resect DNA double-strand break ends. Cell. 134 (2008) 981–994.

16. J. Hu, L. Sun, F. Shen, Y. Chen, Y. Hua, Y. Liu, M. Zhang, Y. Hu, Q. Wang, W. Xu, F. Sun, J. Ji, J. M. Murray, A. M. Carr, D. Kong, The intra-S phase checkpoint targets Dna2 to prevent stalled replication forks from reversing. Cell. 149(6) (2012) 1221–1232. doi: 10.1016/j.cell.2012.04.030.

17. S. Thangavel, M. Berti, M. Levikova, C. Pinto, S. Gomathinayagam, M. Vujanovic, R. Zellweger, H. Moore, E. H. Lee, E. A. Hendrickson, P. Cejka, S. Stewart, M. Lopes, A. Vindigni, DNA2 drives processing and restart of reversed replication forks in human cells. J Cell Biol. 208(5) (2015) 545–562. doi: 10.1083/jcb.201406100.

18. P. Cejka, DNA End Resection: Nucleases Team Up with the Right Partners to Initiate Homologous Recombination. J Biol Chem. 290(38) (2015) 22931–22938. doi: 10.1074/jbc.R115.675942.

19. G. Ölmezer, M. Levikova, D. Klein, B. Falquet, G. A. Fontana, P. Cejka, U. Rass, Replication intermediates that escape Dna2 activity are processed by Holliday junction resolvase Yen1. Nat Commun. 7 (2016) 13157. doi: 10.1038/ncomms13157.

20. R. A. Deshpande, T. E. Wilson, Modes of interaction among yeast Nej1, Lif1 and Dnl4 proteins and comparison to human XLF, XRCC4 and Lig4. DNA Repair (Amst) 6 (2007) 1507–1516.

21. B. L. Mahaney, S. P. Lees-Miller, J. A. Cobb, The C-terminus of Nej1 is critical for nuclear localization and non-homologous end-joining. DNA Repair (Amst). 14 (2014) 9–16. doi: 10.1016/j.dnarep.2013.12.002.

22. K. Yano, K. Morotomi-Yano, K. J. Lee, D. J. Chen, Functional significance of the interaction with Ku in DNA double-strand break recognition of XLF. FEBS Lett. 585(6) (2011) 841–846. doi: 10.1016/j.febslet.2011.02.020.

23. G. Grundy, S. Rulten, R. Arribas-Bosacoma, K. Davidson, Z. Kozik, A. W. Oliver, L. H. Pearl, K. W. Caldecott, The Ku-binding motif is a conserved module for recruitment and stimulation of non-homologous end-joining proteins. Nat Commun. 7 (2016) 11242. https://doi.org/10.1038/ncomms11242

24. C. Nemoz, V. Ropars, P. Frit, A. Gontier, P. Drevet, J. Yu, R. Guerois, A. Pitois, A. Comte, C. Delteil, N. Barboule, P. Legrand, S. Baconnais, Y. Yin, S. Tadi, E. Barbet-Massin, I. Berger, E. Le Cam, M. Modesti, E. Rothenberg, P. Calsou, J. B. Charbonnier, XLF and APLF bind Ku80 at two remote sites to ensure DNA repair by non-homologous end joining. Nat Struct Mol Biol. 25(10) (2018) 971–980. doi: 10.1038/s41594-018-0133-6.

25. M. Sulek, R. Yarrington, G. McGibbon, J. D. Boeke, M. Junop, A critical role for the C-terminus of Nej1 protein in Lif1p association, DNA binding and non-homologous end-joining. DNA Repair (Amst). 6(12) (2007) 1805–1818. doi: 10.1016/j.dnarep.2007.07.009.

26. M. Ferrari, D. Dibitetto, G. De Gregorio, V.V. Eapen, C.C. Rawal, F. Lazzaro, M. Tsabar, F. Marini, J.E. Haber, A. Pellicioli, Functional interplay between the 53BP1-ortholog Rad9 and the Mre11 complex regulates resection, end-tethering and repair of a double-strand break. PLoS Genet. 11 (2015) e1004928. doi: 10.1371/journal.pgen.1004928.

27. J. F. Gilles, M. Dos Santos, T. Boudier, S. Bolte, N. Heck DiAna, an ImageJ tool for object-based 3D co-localization and distance analysis. Methods. 115 (2017) 55–64. doi: 10.1016/j.ymeth.2016.11.016.

28. A. Mojumdar, M. De March, F. Marino, S. Onesti, The Human RecQ4 Helicase Contains a Functional RecQ C-terminal Region (RQC) That Is Essential for Activity. J Biol Chem. 292(10) (2017) 4176–4184. doi:10.1074/jbc.M116.767954

29. S. M. Carney, A. T. Moreno, S. C. Piatt, M. Cisneros-Aguirre, F. W. Lopezcolorado, J. M. Stark, J. J. Loparo, XLF acts as a flexible connector during non-homologous end joining. eLife. 9 (2020) e61920. https://doi.org/10.7554/eLife.61920

30. V. Marini, L. Krejci, Unwinding of synthetic replication and recombination substrates by Srs2. DNA Repair (Amst). 11(10) (2012) 789–798. doi:10.1016/j.dnarep.2012.05.007

31. H. H. Y. Chang, N.R. Pannunzio, N. Adachi, M. R. Lieber, Non-homologous DNA end joining and alternative pathways to double-strand break repair. Nat Rev Mol Cell Biol. 18 (2017) 495–506, doi:10.1038/nrm.2017.48.

32. M. Hohl, A. Mojumdar, S. Hailemariam, V. Kuryavyi, F. Ghisays, K. Sorenson, M. Chang, B.S. Taylor, D.J. Patel, P.M. Burgers, J.A. Cobb, J.H.J. Petrini, Modeling cancer genomic data in yeast reveals selection against ATM function during tumorigenesis. PLoS Genet. 16 (02020) e1008422. doi: 10.1371/journal.pgen.1008422.

33. M. A. Slatter, A. R. Gennery, Update on DNA-Double Strand Break Repair Defects in Combined Primary Immunodeficiency. Curr Allergy Asthma Rep. 20(10) (2020) 57. doi:10.1007/s11882-020-00955-z.

